# Molecular Dynamics-Guided Design and Chemoproteomic Profiling of Covalent Kinase Activity Probes

**DOI:** 10.1101/2025.10.17.683178

**Authors:** Pratyasha Chakraborty, Anthony Carlos, Trayder Thomas, Kyle Ghaby, Shaghayegh Fathi, Mukta Sharma, Ngoc Kim Nguyen, Mason Farmwald, Lydia Blachowicz, Shaopeng Yu, Kavya Smitha Pillai, Benoît Roux, Raymond Moellering

## Abstract

Covalent small molecule activity probes can be powerful tools to interrogate protein function in native cellular environments. The design of family-wide activity probes requires an understanding of the molecular sources of general targeting potential and specificity to enable broad targeting of protein family members. Here, we developed and applied a multifaceted docking and molecular dynamics (MD) simulation pipeline to design and test cell-permeable covalent kinase activity probes from a set of hinge-binding pharmacophores. This computationally-guided approach yielded a new cell-active probe, K60P, which targets around 114 kinases across distinct kinase classes in live cells. Chemoproteomic profiling of this probe and a clinical candidate sharing the same indazole core, KW-2449, identified kinase and non-kinase target profiles that differ from recombinant protein assay profiles, underscoring the utility of native kinase profiling *in situ*. Biochemical studies with a model target kinase, ABL1, confirmed covalent labeling of the active site lysine across several kinase probes with distinct kinetics, as well as covalent labeling of key tyrosines *in trans* between ABL1 monomers. Finally, focused proteomics, kinetic modeling, and molecular dynamics simulations revealed that K60P, as well as the comparator probe XO44, preferentially engage with target kinases in their active, DFG-in conformations, which is driven by increasing population of reaction-ready small molecule conformation. These results together establish a computational and kinetic modeling framework for designing covalent activity probes and highlight the balance of target selectivity and kinetic efficiency as a key factor in determining their proteome-wide reactivity.

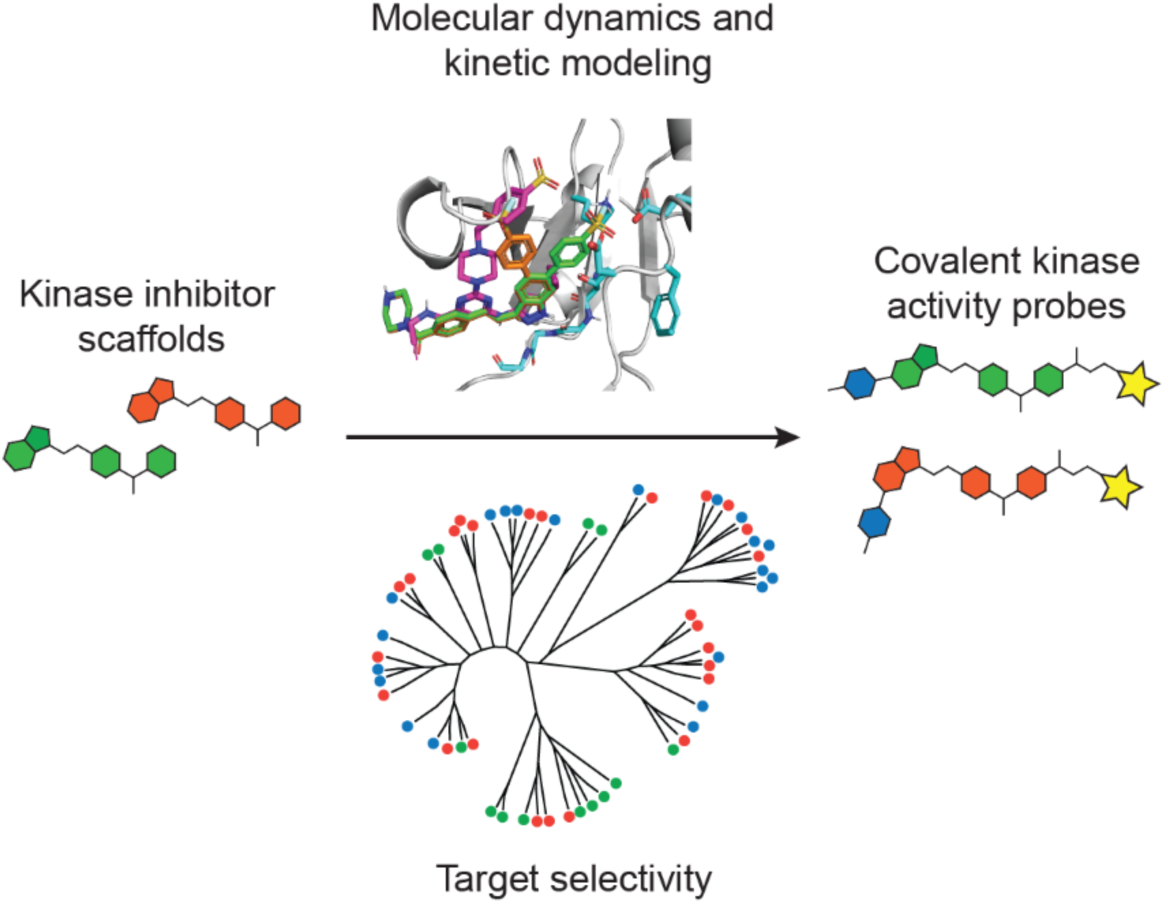

## Introduction

Protein kinases are involved in regulation of diverse cellular processes, including metabolism, cell cycle control, transcription, cell migration and cell death. Aberrant kinase activity contributes to many diseases, most notably cancers, where mutation, translocation, or amplification of specific kinases can contribute to unchecked proliferative signaling. Advances in medicinal chemistry and structure-based design have resulted in the development of small molecule inhibitors and FDA-approved drugs targeting conserved ATP-binding active sites of dozens of kinases for various disease indications. However, the fine-tuning of potent activity over structurally related kinases is an important aspect of kinase inhibitor development. In particular, identification of the on-and off-target interaction across the kinome as well as the broader proteome remains a challenge, and many kinase inhibitors are known to exhibit polypharmacology^1–6^. The disparity between recombinant assays compared to that against native kinase targets in cells is well known. Technologies like KinomeScan^4, 5^, Kinobeads^6^, and KiNativ^7^ have been widely deployed to identify the target landscape of kinase inhibitors using either recombinant proteins or native proteins in cell lysates. However, these approaches suffer from several drawbacks. For instance, KinomeScan requires the genetic engineering or purification of recombinant proteins, typically just interrogating the catalytic domain of target kinases and thereby does not replicate native cellular conditions. Kinobeads utilize immobilized kinase inhibitors to scan for native protein kinase binding in cell lysates, and KiNativ employs a reactive ATP-mimetic probe to label active kinases; both technologies perform these measurements in the artificial cell lysate environment. There remains a need for *in situ* profiling of native kinases and streamlined proteomic workflows to profile small molecule probes and potential selective therapeutics.

Covalent small molecules are attractive as both probes and therapeutics due to their potential for increased potencies, improved target residence kinetics, and high selectivity. Recently, several cysteine-targeted covalent drugs have been approved in cancer, including molecules targeting wild-type cysteines adjacent to the ATP binding site in protein kinases^8, 9^ or neomorphic mutant cysteines in GTPases like k-RAS^10^. Targeting non-catalytic cysteines near the ATP-binding site has served as a general approach to design irreversible kinase probes, including several FDA-approved drugs^11, 12^. Expansion to other nucleophiles like lysine and tyrosine is increasingly gaining popularity to expand the targetable kinome due to the absence of druggable cysteines near the active site of many kinases. Activated esters, vinyl sulfones, sulfonamides, and sulfonyl fluorides have been used to target lysines^13^. In particular, sulfonyl fluorides have been widely used to target lysines due to their desired reactivity and hydrolytic stability in biological environments. For example, Taunton and coworkers installed a sulfonyl fluoride warhead on pyrimidine 3-aminopyrazole, which is a promiscuous kinase-binding scaffold, to generate a general lysine-reactive kinase probe (XO44) that can label active kinases directly in cells. XO44 and its fluorosulfonate and aldehyde analogs have been used to profile the catalytic lysines of the kinome^14, 15^, permitting interrogation of hundreds of kinases. However, XO44 targets a subset of the kinome and requires downstream click-chemistry labeling for target detection and quantification, potentially limiting applications like spatial or low-sample-input profiling^16–20^. Additionally, XO44 also labels many other proteins owing to its highly reactive sulfonyl fluoride covalent warhead, raising the question of whether less reactive kinase probes could be useful for in-cell or live tissue profiling studies.

Here, we elaborate a virtual screening workflow that combines docking and molecular dynamics (MD) to develop and study several covalent kinase activity probe scaffolds. This work follows considerable recent interest in applying and expanding computational screening methods for the design of covalent ligands^21, 22^. Guided by the computational modeling, we considered several core kinase-targeting scaffolds and ultimately synthesized and tested a novel sulfonylfluoride-modified indazole probe, K60P, that targets the catalytic lysine of kinases, specifically in their active conformations, *in vitro* and in cells. We also synthesized a biotinylated XO44 analog, X4K, to enable click-free in-cell protein labeling and streamline readouts and included it as a comparator in our proteomics and biochemical assays. Our integrated computational, biochemical, and cell-based profiling workflow helped rationalize kinetic determinants of selectivity and potency for specific kinases, including ABL1 as a model kinase.

## Results

### Computational Design of covalent kinase activity probes identifies K60P as candidate

When considering core kinase-targeting for the design of cell permeable activity probes, we focused on several hinge-binding scaffolds with reasonably promiscuous *in vitro* kinase profiles. These included amino-pyrazoles, pyridyl-imidazoles, indazoles and others, which are found in pre-clinical and clinical kinase inhibitors (e.g., sunitinib^23^ and KW-2449^24^). To identify permissive sites for covalent warhead introduction around these hinge-binding core scaffolds, we developed a computational workflow comprised of docking followed by MD simulations. From initial studies and published in vitro kinome profiles^4, 5^, we determined that the compact indazole pharmacophore related to the phase 1 clinical candidate KW-2449 was a suitable starting point for development of pan-or multi-kinase covalent probes. We determined the analog space to screen based on inspection of the predicted bound pose of KW-2449, inferred from its shared scaffold with axitinib, which has been co-crystallized with ABL1^25^. We also aligned representative members from seven kinase groups against the ABL1/KW-2449 model (Figure 1A, Figure S1A) which revealed conserved alignment of hinge-binding residues and proximity of conserved catalytic lysines to the indazole and amino pyrimidine cores of KW-2449 and XO44, respectively. The alignment also highlighted the need to stay clear of the gatekeeper residue to avoid individual kinase selectivity^26^.

**Figure 1:**
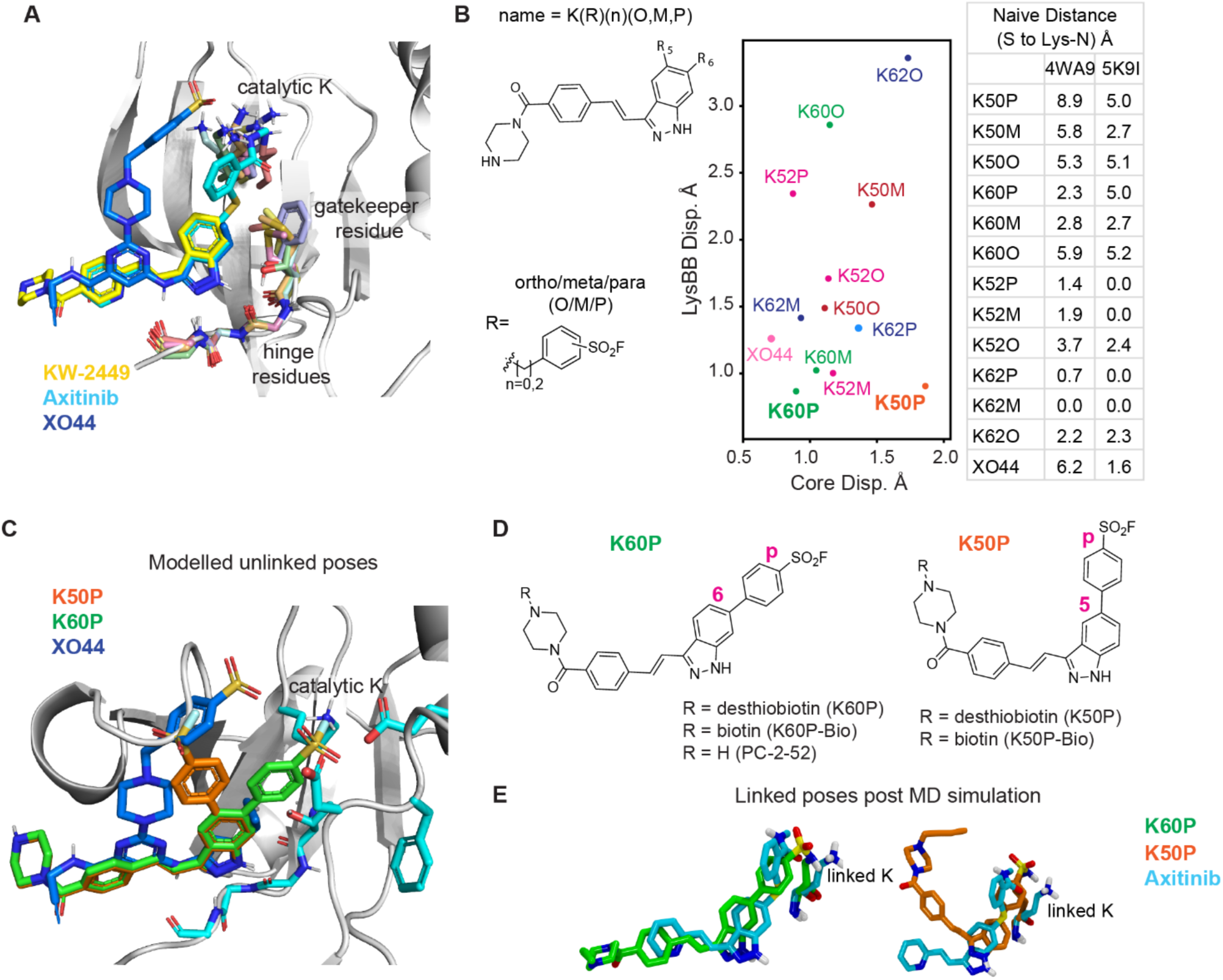
Computation-guided design for covalent, conserved lysine-targeting kinase probes. (**A**) Overlayed binding poses of indicated ligands within the conserved ATP-binding pocket defined by features shared between representative kinases CAMK1D, AKT1, MAP2K1, ALK2, ABL1, CDK1, CK1α. (**B**) Left: probe library scaffold structure and naming convention. Middle: lysine backbone and core displacements (in Å) by each probe analog. Right: minimum achievable distance between the SO_2_F warhead and the crystallographic pose of the lysine in 4WA9 (ABL1) and 5K9I (SRC) for each anchored analog. (**C**) Modeled unlinked poses of K60P (green), K50P (orange), and XO44 (blue) in ABL1 kinase, illustrating warhead positioning. (**D**) Chemical structures of K60P and K50P. (**E**) Final linked poses of K60P (left, green), K50P (right, orange), and linked lysine (yellow) after Molecular Dynamics (MD) simulations (100 ns). The crystallographic pose of Axitinib (cyan) and lysine is shown for comparison.

We focused on a library of analogs containing a phenylsulfonyl fluoride group introduced at the 4, 5, or 6 positions of the indazole heterocycle connected with linker lengths of 0, 1, or 2 atoms (Figure 1B, Figure S1B). Initial docking was performed as a coarse screen to rapidly eliminate analogs where the addition of the linker and phenyl group introduced steric clashes or positioned the warhead outside of the binding pocket. The receptors for docking were 4WA9 (ABL1 + axitinib), 5K9I (SRC + XO44), and 3G0E (KIT + sunitinib), and were chosen based on ligands of interest and to expand the binding cavity around potential scaffold modification positions. Analogs that docked in multiple viable poses across multiple receptors were considered to have the highest potential. From this evaluation, analogs modified at the 5 or 6 position of the indazole, with linker lengths of 0 or 2 atoms were identified as the most promising cores for further study. We also included designs with a retrievable desthiobiotin or biotin tag linked to the piperazine of the KW-2449 pharmacophore for further computational screening, since neither tag interfered with target binding.

We subjected 12 analogs to further scrutiny by conducting unbiased MD simulations of their covalently linked complex, further expanding the set of analogs to account for the possibility of attaching the warhead at the ortho-, meta-, or para-positions of the phenyl ring. Each analog was initially aligned to the crystallographic pose of axitinib in ABL1 kinase (PDB ID: 4WA9, chain B). A covalent bond was then introduced in the force field between the conserved Lys271 in ABL1 and the electrophilic warhead sulfur for each analog. The systems were equilibrated with the ligand’s indazole nitrogens and protein restrained, prior to being released for 100 ns of unbiased MD simulation with explicit solvent. The structural strain of the covalently linked pose was evaluated over the final 10 ns of MD as a combination of two metrics: 1) the position of the ligand indazole core relative to the axitinib defined pose; 2) any perturbation of the lysine backbone relative to the protein crystal structure (Figure 1B). We assumed that both metrics are reporters of the same strain within the covalently linked structure, with a high score in either indicating that the ligand in the kinase active site would be unlikely to adopt a suitable reaction-ready conformation with the catalytic lysine before covalent engagement. We also scored the candidates based on the minimum warhead-S to lysine-N distance between a conformational search of the linker and warhead of each analog and to the crystallographic lysine-N positions of structures 4WA9 and 5K9I with no consideration for sterics as a “naïve” structural metric (Figure 1B table). With the exception of K50P and K62O, all ligands maintained key hydrogen bond interactions to the hinge, although many did so from more strained conformations (Figure S1C). Though several promising scaffolds were found, K60P was selected for synthesis based on the metrics as the best performing analog overall (Figure 1C-E). We also synthesized a less promising analog K50P, which showed a less favorable core displacement after bond formation with Lys271, but was still predicted to bind reversibly (Figure 1C-E, Figure S2).

### Proteome-wide profiling reveals kinase and non-kinase targets of K60P and KW-2449

We next explored the utility of K50P and K60P as cell-permeable kinase activity probes by probing the proteome-wide labeling of their desthiobiotin-labeled versions. Both probes showed dose-dependent covalent labeling of endogenous protein targets, as visualized by streptavidin immunoblotting of desthiobiotin-tagged proteins in cell lysates and intact cells (Figure 2A). These data confirmed the suitable stability and cell permeability for *in-situ* profiling of these probes. In accordance with our modeling predictions, K60P exhibited more robust labeling of the proteome than K50P in western blot-based assays. We therefore proceeded with K60P for further downstream analyses to identify the probe-labeled proteins. To identify the targets of K60P quantitatively, we synthesized the biotin-labeled version of K60P, K60P-Bio, and performed LC-MS/MS analysis using stable isotope labeling by amino acids in cell culture (SILAC) for quantitation of probe-labeled proteins (Figure 2B). We treated K562, HeLa, and MCF7 cell lines with K60P-Bio, enriching probe-labeled proteins on streptavidin beads. We considered target proteins as ‘enriched’ if they exhibited consistent SILAC ratios of at least 2 in probe-treated, ‘heavy’ cells relative to vehicle-treated, ‘light’ cells after processing and LC-MS/MS analysis. K60P-Bio enriched a total of 114 protein kinases in these three cell lines combined (Figure 2C-D, Table S1). Among these enriched protein kinases, we identified known targets of the parent scaffold KW-2449: ABL1, AURKA and B, PAK2, BMP2K, RIPK1, TNIK, DYRK1A, and CSNK2A1 and A2^5, 6, 27^. We also observed enrichment of kinases not previously annotated as targets of KW-2449, including YES1, LIMK1, TAOK1, 2, 3, CDK1, 2, 4, 5, 6, and RSK1 and 2. Several metabolic kinases, including HK1 and 2, DTYMK, AGK, ADK, CKB, and PANK4 were also enriched. Intriguingly, the kinome coverage of K60P was reduced compared to a biotinylated version of XO44, X4K, that we synthesized and profiled in K562 cells (Figure 2C, Figure S3, Table S2). X4K enriched roughly twice as many kinases in this context alongside a much larger total number of targets (∼1800), which could be related to higher intrinsic reactivity in the proteome (*vide infra*).

**Figure 2:**
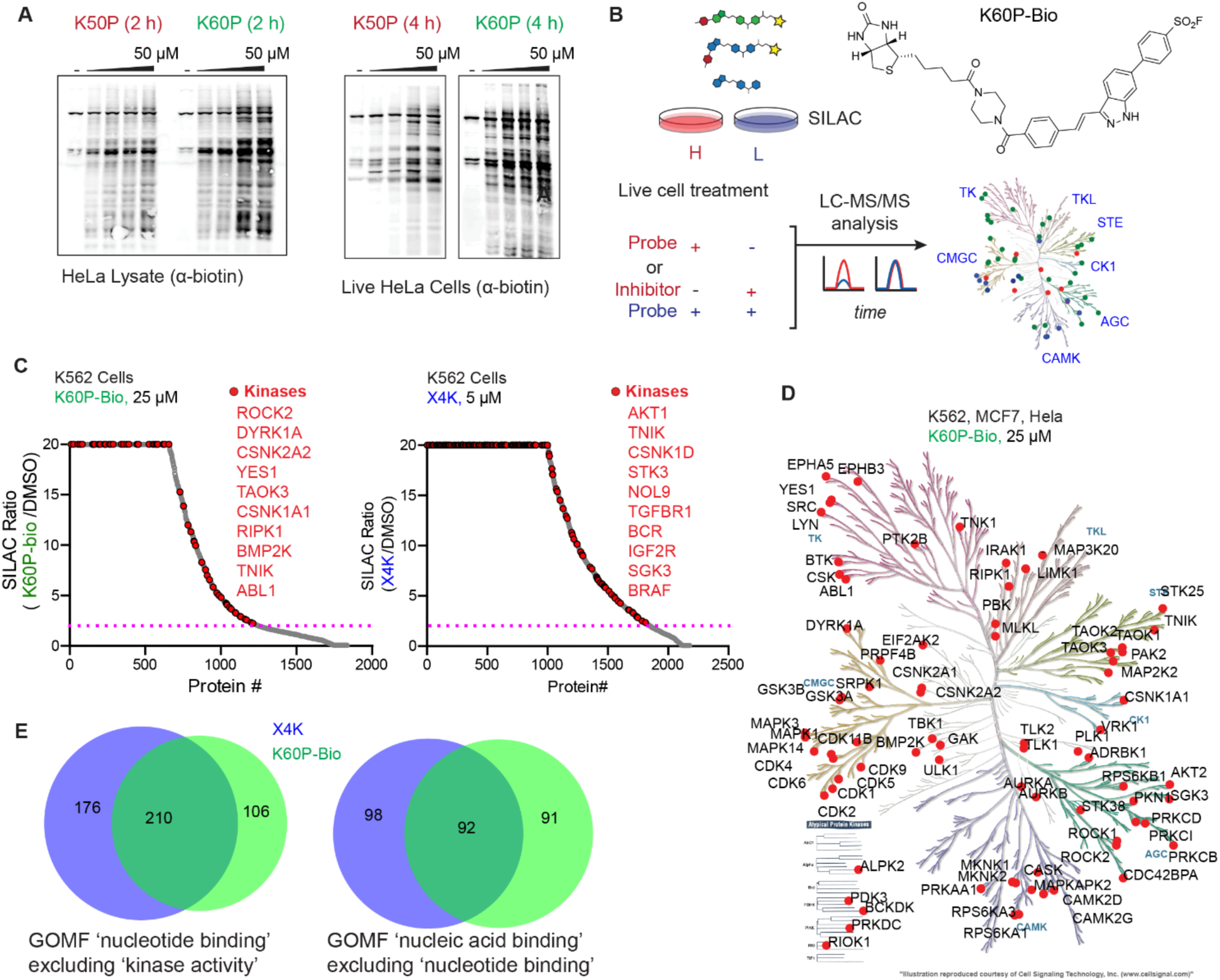
Proteome-wide labeling by K60P-Bio in lysates and live cells. (**A**) Representative Western blots demonstrate broad labeling of proteins in lysates (left) and in live cells (right) by K60P and K50P. (**B**) Workflow of kinase probe SILAC-MS/MS profiling experiments to identify proteins labeled by K60P-Bio (structure shown). (**C**) Waterfall plots of kinases labeled by K60P-Bio or X4K in live K562 cells treated with either K60P-Bio (25 µM for 4 hours, left) or X4K (5 µM for 1 hour, right). Control cells were treated with DMSO. Cutoff: SILAC Ratio ≧ 2 is shown as dotted line; n = 2 (**D**) Dendogram showing kinases enriched by K60P-Bio in K562, Hela, and MCF7 cell lines, n = 2 for each cell line. (**E**) Venn diagram of shared and unique nucleotide-and nucleic acid-binding proteins labeled by K60P-Bio and X4K in live K562 cells.

K60P is closely related to the parent indazole, KW-2449, which was assessed in the clinic for targeting FLT3 and potentially ABL1, two tyrosine kinases. Despite this structural origin, K60P demonstrated less engagement of TK and TKL kinase sub-families relative to X4K. Common targets between the two probes in these families included YES, LYN, BTK, CSK, PYK2, LIMK, RIPK1 and ABL1. By contrast, K60P engaged considerably more kinases in the STE, CK, AGC and CAMK sub-families, which extends the coverage of the kinome across these two cell permeable probes (Figure 2D). Beyond kinases, approximately 30% of the proteins labeled by both probes were annotated as ‘nucleotide binding’ from gene ontology analyses (Figure 2E). Additional classes of K60P-Bio and X4K-enriched proteins included those involved in protein synthesis, nucleic acid binding, and cytoskeletal proteins (Figure 2E, Figure S4). This likely suggests that these off-targets are highly abundant proteins, rich in solvent-exposed lysines and tyrosines, which are non-specifically labeled by the electrophilic sulfonyl fluoride warhead, regardless of the probe scaffold^28^.

Because K60P engaged a distinct subset of kinase and non-kinase targets in cells, we sought to use it in a competitive format to identify targets of the reversible parent scaffold KW-2449. SILAC-labeled K562 cells were pre-treated with 2 µM KW-2449 or DMSO for 1 hour prior to K60P-Bio treatment, target enrichment, and analysis by LC-MS/MS. NTRK and TNIK^6, 29^, which are known targets of KW-2449, were significantly competed under these conditions. Several novel kinase targets, including RPS6KB2, MAP3K20, GSK3A, VRK2, as well as the metabolic kinases AGK and HK2, were significantly competed (Figure 3A-B, Table S3). These data suggest that KW-2449 may exert considerable background target engagement of housekeeping enzymes under physiologically relevant conditions. We note that competition between a covalent probe and a non-covalent competitor biases towards capture of higher affinity targets and may miss other transient interaction partners. Metascape analysis of competed KW-2449 targets indicated strong enrichment of proteins with reported localization to mitochondria (GO Cellular Component), including several kinases (HK2, RAF1, VRK2 and AGK, Figure 3C), which could be related to the piperazine linker in this molecule and many other small molecule kinase inhibitors. There were numerous non-kinase targets that were significantly competed by KW-2449, about 30% of these included several GDP/GTP binding proteins such as DNM1, RAB proteins, and the immunomodulatory protein STING1 (Figure 3D). This could potentially be attributed to binding site structural similarity, perhaps indicating a degree of molecular crosstalk that would be missed in recombinant kinase assays. Notably, ABL1 was not significantly competed by KW-2449, despite it being previously reported as a high-affinity and -value clinical target for development^29^. This conclusion agrees with published bead-based profiling of native proteins in lysates^6^ but runs counter to work showing that KW-2449 potently targets the catalytic domain of ABL1 and other kinases^4^.

**Figure 3:**
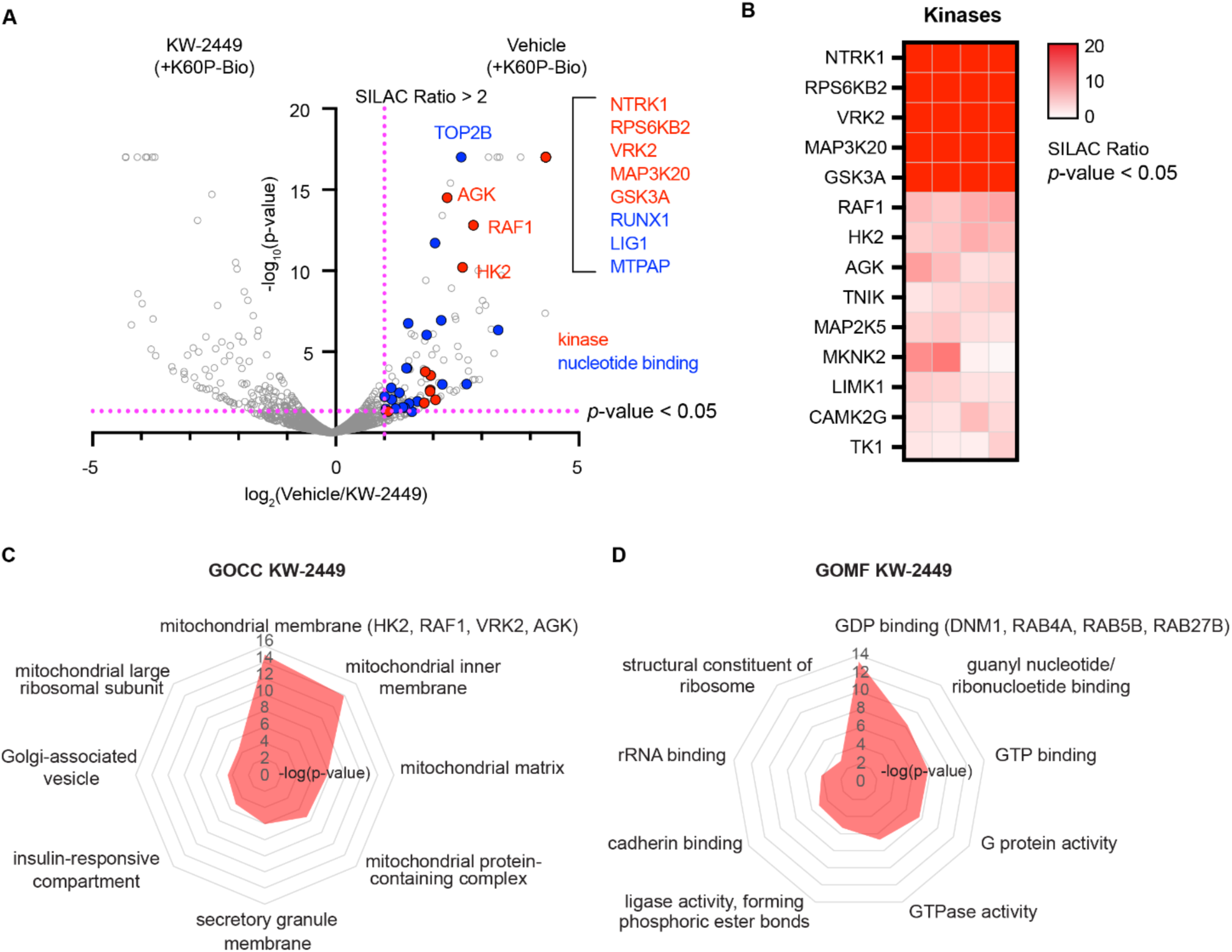
Competition with K60P-Bio reveals known and novel targets of KW-2449 in live cells. **(A)** Volcano plot of streptavidin-enriched SILAC ratios for K562 cells treated with 2 µM KW-2449 (1 hour) followed by with 25 µM K60P-Bio (2 hours). Cutoffs: competition SILAC Ratio ≧ 2, *p* < 0.05; n = 4. (**B**) Heatmap of kinases competed by KW-2449 by over 2-fold. (**C-D**) Gene ontology cellular component (C) and molecular function (D) bioinformatic enrichment analysis of all protein targets significantly competed by KW-2449 (*p* < 0.05).

### K60P Can Covalently Label the Active Conformation of ABL1 Through both Cis and Trans Covalent Labeling Mechanisms

To understand the mechanisms underlying active site engagement and covalent modification by K60P and close analogs, we performed biochemical and biophysical studies with recombinant ABL1, a reported target of the progenitor scaffold and a focus of our computational screening effort. We performed dose-and time-dependent labeling studies with recombinant ABL1 (residues 46-515) and K60P, K50P, and X4K. Western blot analyses confirmed sub-micromolar covalent labeling of ABL1 by K60P (EC_50_ = 0.6 µM) and X4K (EC_50_ = 0.7 µM), and less potent engagement by K50P (EC_50_ = 2.9 µM) (Figure 4A, Figure S5). Preincubation with a non-desthiobiotinylated analog of K60P (PC-2-52) resulted in complete competition with both K60P and X4K with sub-micromolar IC_50_ values (0.5-1 µM), suggesting that the predominant covalent labeling was occurring within the active site of ABL1 (Figure 4B, Figure S6). Time-dependent studies showed X4K labeled recombinant protein to apparent saturation within minutes, which was much faster than either K60P or K50P (Figure 4C, Figure S5). These results were surprising given the similar arylsulfonylfluoride warhead and molecular dynamics scores between K60P and XO44 for ABL1. This kinetic advantage could contribute to the apparent increase in kinome labeling by X4K, as well as higher labeling of non-kinase targets in the proteome.

**Figure 4:**
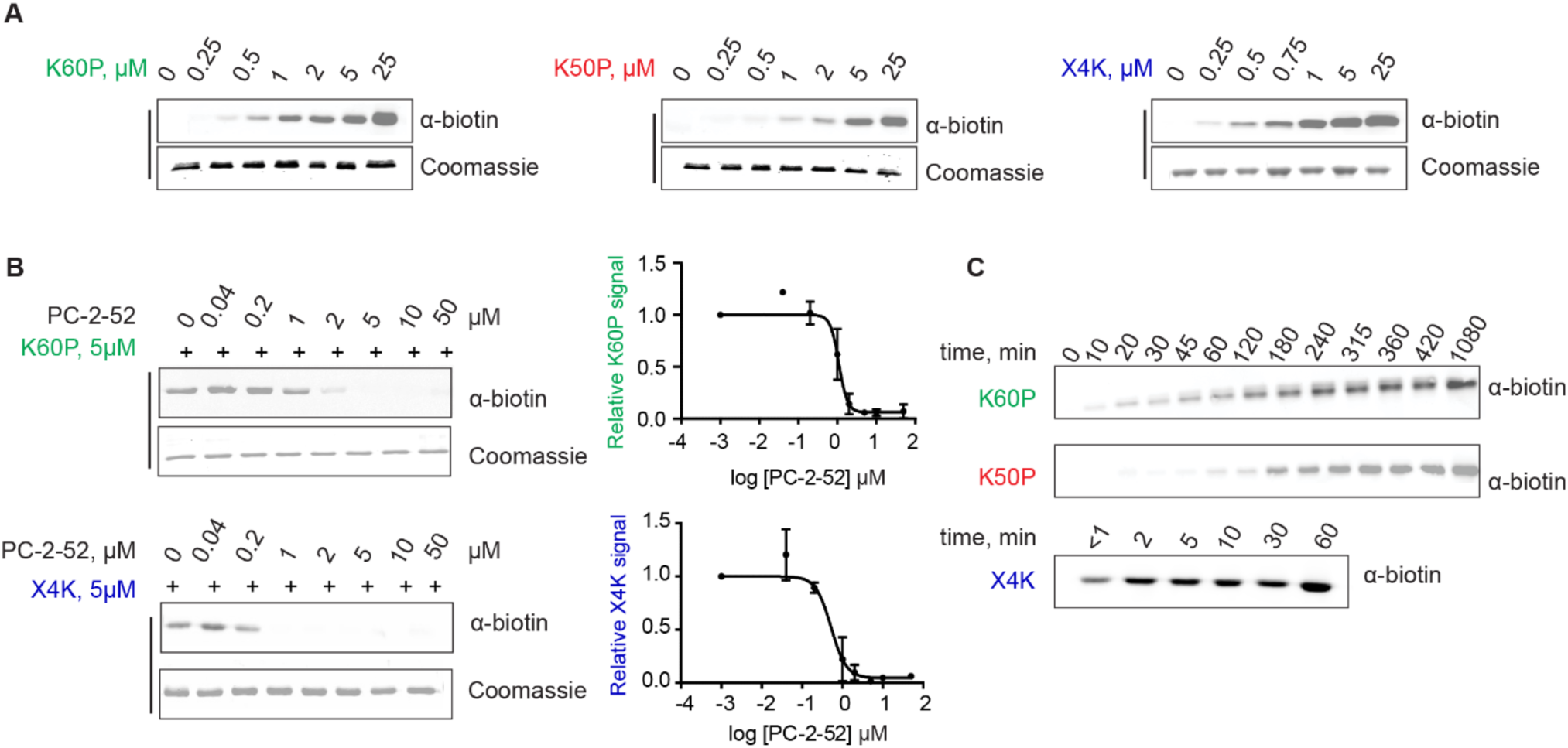
K60P, K50P, and X4K covalently label recombinant ABL1 with distinct kinetic profiles. (**A**) Streptavidin immunoblot showing dose-dependent labeling of the recombinant ABL1 protein with K60P (left), K50P (middle), and X4K (right) (0-25 µM, 1 hour), n = 2. Coomassie stain was used as loading control. (**B**) Preincubation of recombinant ABL1 with PC-2-52 (non-biotinylated analog of K60P) dose-dependently decreases recombinant ABL1 labeling by both K60P (top) and X4K (bottom), n = 2. (**C**) Streptavidin immunoblot showing time-dependent labeling of recombinant ABL1 with K60P and K50P (10 µM, up to 18 hours) and X4K (10 µM, up to 1 hour), n = 2.

To confirm and compare the site(s) of covalent labeling by K60P and X4K, we performed focused proteomic mapping to detect and quantify the covalent adducts between each probe across the recombinant ABL1 surface. Recombinant ABL1 was incubated with each probe at 10 µM for 1 hour, followed by digestion and processing for peptide-level detection of covalent adducts across lysine and tyrosine residues in the protein. We observed direct modification of the catalytic target lysine, K271, as the dominant adduct by both peptide spectral matches (PSMs) and ion intensity for both K60P and X4K (Figure 5A-B; Figure S7A). Two other sites of modification were also observed for K60P: Y226, located in the SH2 kinase linker, and Y393 in the activation loop (A loop), though these peptides were observed at lower relative ion intensities (Figure 5B-C, Figure S7B). By contrast, X4K labeling was observed at seven unique sites within ABL1, including the aforementioned K271, Y226, and Y393, as well as Y115, Y215, Y139, and K262 (Figure 5B-C, Figure S7B). Inspection of available ABL1 structures indicated that the phenolic oxygens in Y226 and Y393 do not come into reasonable proximity with the bound sulfonylfluoride warhead, however two possible configurations with distance of less than 5 Å between Tyr393 and the K60P warhead location could be found within a 400 µs MD dataset of apo-ABL1 seeded from ABL1 crystal structures that was previously reported by Meng *et al*. (Figure 5C, Figure S7C)^30^. Sulfonylfluorides react more readily with tyrosine than lysine^31^ but collectively this data suggests that such poses are short-lived and relatively rare. Another possibility supported by this data involves covalent labeling of the observed tyrosines *in trans* on another bound-ABL1 monomer. These sites are the primary sites of *trans* auto-phosphorylation^32, 33^, providing a tantalizing possibility that the phosphorylatable tyrosines of an active kinase monomer could be labeled by a probe reversibly bound to a second kinase monomer, through a similar mechanism (Figure 5D). Indeed, reversibly bound K60P places the reactive warhead in proximity of the catalytic lysine as well as in the vicinity of the active site entrance involved in phosphotransfer to these tyrosine targets of auto-phosphorylation (Figure 5E). This mechanism would support the possibility of covalent targeting of kinase substrates *in trans* (Figure 5D-E); additional focused work to elucidate the mechanism and generality underlying these observations is needed.

**Figure 5:**
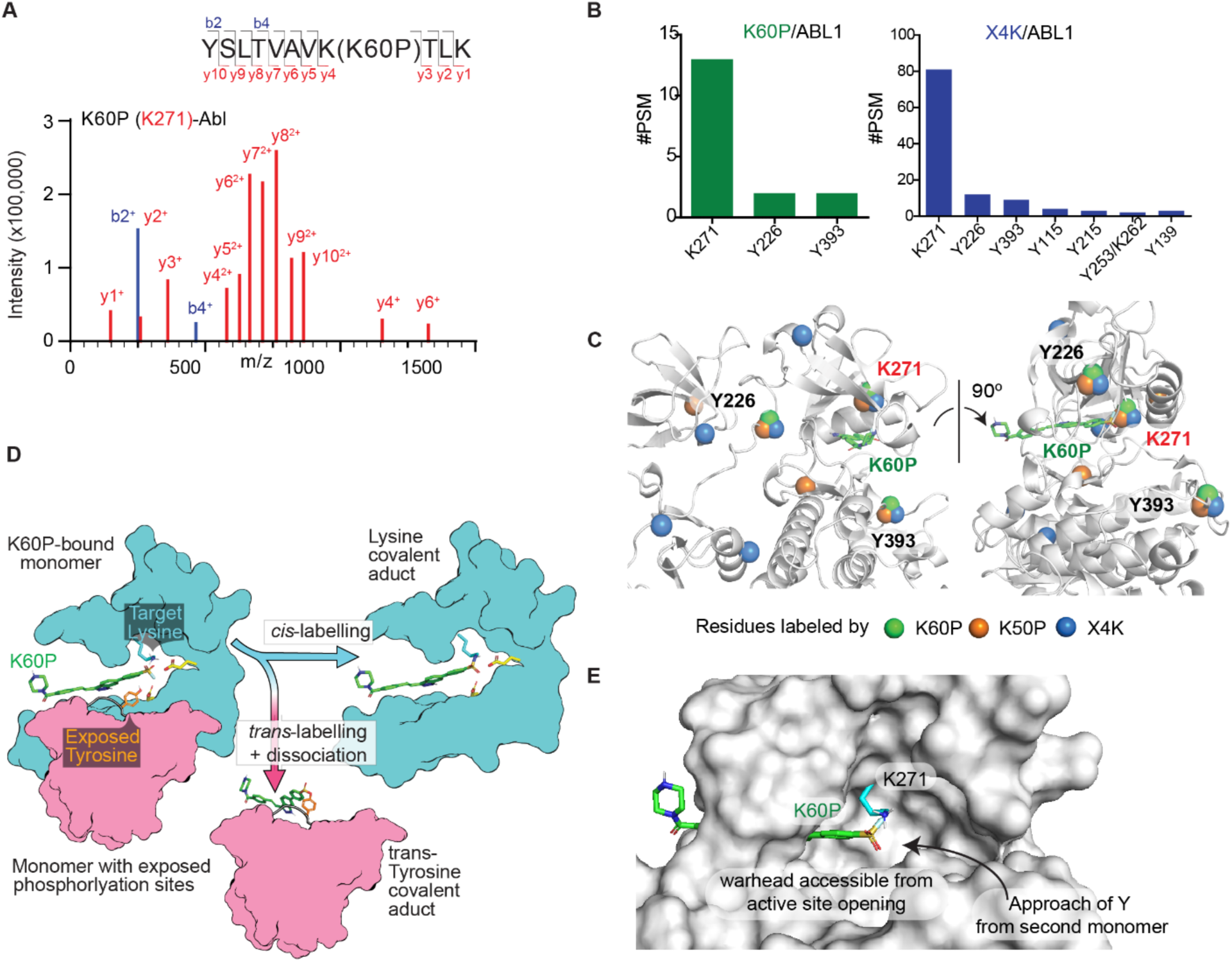
K60P and X4K covalently label distinct sites in ABL1 with *cis*-labeling of conserved lysine and a proposed *trans*-labeling of phosphorylatable tyrosines. (**A**) MS/MS spectrum of trypsinized ABL1/K60P adduct showing the labeled K271. (**B**) ABL1 sites labeled by K60P (green, left) and X4K (blue, right). Bar graphs show the number of peptide spectral matches (#PSMs) for each site. (**C**) ABL1 structure showing the positions of residues labeled by K60P (green), K50P (orange), and X4K (blue) relative to K60P docked in the active site (green). K271, Y226, and Y393 are labeled by all probes. (**D**) Proposed model for *cis*-labeling of K271 and *trans*-labeling of Y226/Y393 in ABL1 by K60P. Structural model showing K60P docked in the active site of one ABL1 monomer (teal), either reacting with K271 of the same monomer (*cis*) or with exposed tyrosine phosphorylation sites Y226/Y393 on a neighboring monomer (pink, *trans*). (**E**) K60P docked into ABL1 active site showing that the probe is positioned to access K271 within the same monomer, while its reactive group remains exposed at the active site opening for approach of neighboring monomer tyrosines.

### A Kinetic Modeling Framework to Determine the Contribution of Encounter Complex and Covalent Bond Formation to Probe Labeling Efficiency

Given the disparate kinetics of covalent and kinome-wide labeling of these model sulfonylfluoride probes, as well as the large differences in probe performance in recombinant assays, we sought to rationalize probe performance by quantitatively characterizing the kinetics underlying probe reactivity with ABL1 and other kinases. To do so, we explored a wide range of possible kinetic parameters to fit experimental covalent labeling data within a Bayesian statistical modeling framework using extensive Metropolis Monte-Carlo sampling^34^. In this BMMC procedure, experimental time-and dose-dependent probe labeling data with recombinant ABL1 were fit using parameterized mass-action simulations of irreversible binding between protein P and ligand L (Scheme 1)^35^.

**Scheme 1.**
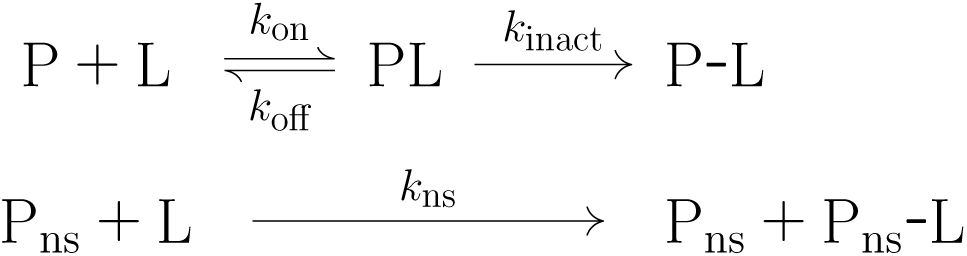
Schematic basis of numerical kinetic simulations.

An initial least-squares (LS) fit normalized the experimental labeling intensities to the fraction of covalently-bound protein and provided a starting point for the BMMC. Sampled parameters included the apparent rate constants of association (*k*_on_), dissociation (*k*_off_), inactivation (*k*_inact_), and nonspecific-binding (*k*_ns_, e.g., non-specific labeling of surface residues). To focus the sampling on the equilibrium binding affinity (*K*_d_=*k*_off_/*k*_on_), for K60P and K50P, the *k*_on_ was fixed (diffusion-limited) while for X4K, *k*_off_ was fixed since *k*_on_ appeared to be rate-limiting. The median (or fixed value) of each parameter together with the 95% highest density interval is tabulated in Table S4. To relate the fitted rate constants with familiar empirical constants of irreversible kinetics, the affinity, the inactivation constant (*K*_I_=(*k*_off_+*k*_inact_)/*k*_on_), and the covalent efficiency (C_eff_=*k*_inact_/*K*_I_) were calculated for each BMMC sample and are also included in the table^36–38^. Simulated dose-and time-responses using reasonable sets of sampled rate constants are shown in Figure 6. The predicted rate constants in the regime of the K50P and K60P indicate a relatively slow *k*_inact_ (Figure S8-10), and thus the rate of inactivation, (i.e. *k*_inact_ ×[PL]), depends on both *k*_inact_ and the affinity (*K*_d_=*k*_off_/*k*_on_). As such, K50P and K60P maintain an effective equilibrium of the reversible binding step and fall in line with the rapid-equilibrium approximation that is fundamental to conventional empirical models^39^. In contrast, the BMMC sampling suggests that X4K displays a relatively fast *k*_inact_, which precludes the equilibrium of the reversible bound state. For X4K, the reversible binding affinity is not resolved by the measurements as the inactivation occurred on a timescale comparable to or faster than reversible binding. Furthermore, the dose-response data for X4K does not appear to reflect a saturation curve of the reversibly associated state, PL, which hinders the precision of rate constant estimations and is explained further in the Supplemental Kinetic Modeling Discussion. Notwithstanding the uncertainty, the modeled lower bound of sampled *k*_inact_ for X4K is over 20 times greater than the upper bound for K60P and the inherent affinity, as fit by *k*_off_/*k*_on_ parameters, does not explain the observed differences. The estimated potency (i.e. covalent efficiency), which describes the proportionality of the rate constants that was conserved across variations of K50P and regimes of X4K, provides a more reliable comparison between the probes, showing that X4K has the highest potency (C_eff_ ≈ 3.3-13.5 × 10⁻⁴ µM⁻¹s⁻¹), followed by K60P (≈ 0.69-2.9 × 10⁻⁴ µM⁻¹s⁻¹) and then K50P (≈ 0.16-0.53 × 10⁻⁴ µM⁻¹s⁻¹). Collectively, these kinetic modeling studies indicate that K60P and K50P likely perform better than X4K in terms of their reversible association, and that the differences in labelling kinetics are driven by *k*_inact_.

**Figure 6:**
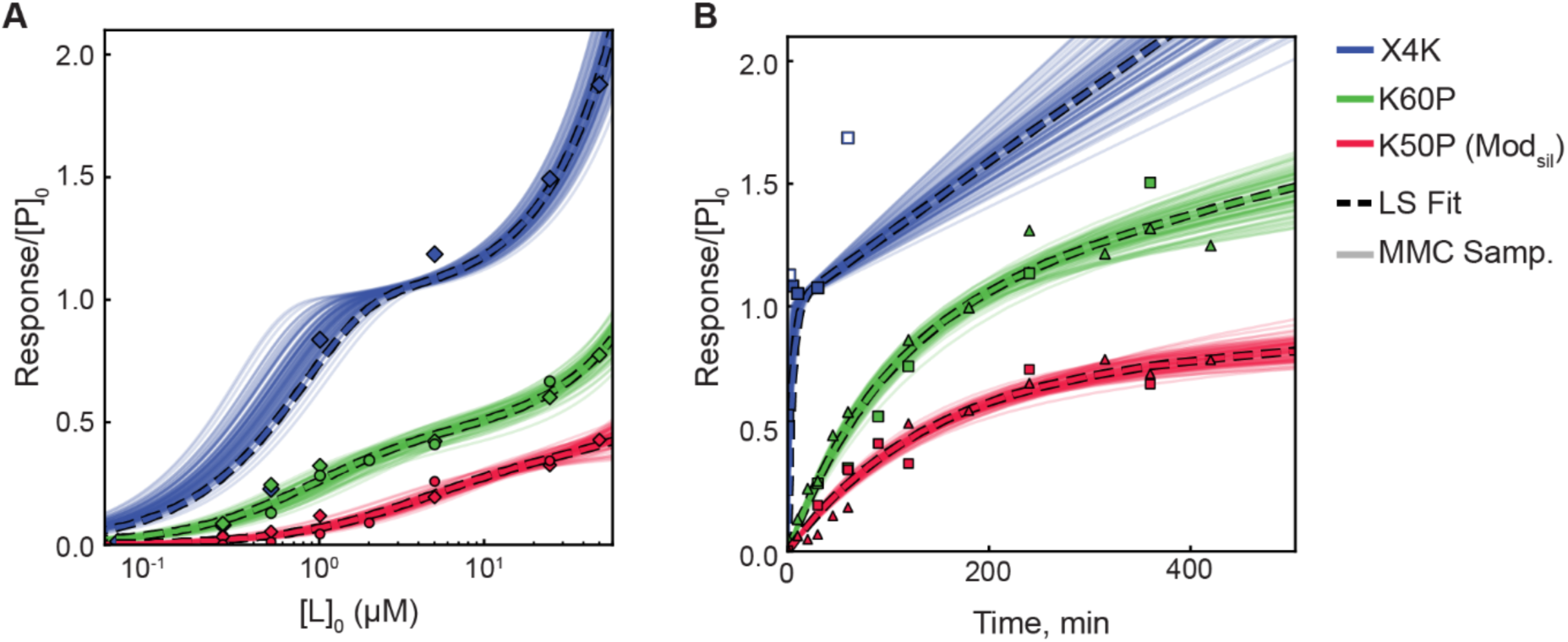
Kinetic modeling reveals complex kinetics of ABL1 labeling by K60P, K50P, and X4K. Experimental and simulated dose-responses (**A**) and time-responses (**B**) of K60P (green), K50P (red), and X4K (blue) binding to ABL1. The experimental data are scaled according to the least-squares (LS) minimization (dashed lines), and the families of solid lines correspond to the kinetic simulations of every 10 BMMC samples whose rate constants exist within their respective 95% highest density intervals. Simulated response is the sum of the specifically-and nonspecifically-bound covalent complexes.

### Molecular Dynamics Confirms Preference for ‘Active’ Kinase Conformations and Rationalizes Inherent Differences in Kinetic Efficiency

Kinetic modelling indicated that the slower rate-limiting processes for K60P and K50P compared to X4K likely occur post-association within the kinase active site. To account for the differences in *k*_inact_, we considered two factors: 1) the differences in the intrinsic reactivity of the warhead leading to the formation of the covalent bond; 2) the likelihood of a reaction-ready arrangement within the binding pocket, in which the target lysine is deprotonated and the ligand warhead is in close-proximity. To assess the intrinsic chemical reactivity of the different warheads to form the covalent bond with the lysine, we conducted labeling experiments of X4K and our probes with bovine serum albumin (BSA) as a model non-specific target protein. This experiment indicated ∼20-fold higher labeling intensity for X4K compared to K60P over the same treatment time (Figure S11). This was higher than, but qualitatively consistent with, the non-specific labeling rates estimated from the kinetic modeling (Table S4). However, Gilbert et al.^31^ explored the kinetics of a series of XO44 analogs and showed that a <100-fold difference in intrinsic reactivity did not correlate directly with significant changes in on-target labeling in model systems. Based on these observations, we conclude that differences in intrinsic reactivity are likely to partly account for the slower rate of labeling for K60P relative to X4K, and intrinsic reactivity of the latter likely contributes to off-target binding profile within the proteome.

Another requirement of the reaction-ready arrangement is the deprotonation of the lysine that enables a nucleophilic attack on the warhead. The likelihood of the lysine being neutral is expected to be influenced by its potential salt-bridges to both the DFG-motif-Asp and αC-helix-Glu. The presence of these salt-bridges has been predicted to result in an increase of two p*K*_a_ units^40^, which is enough to decrease the likelihood of a reaction-ready configuration with a neutral lysine by a factor of 100, and hence alter the apparent rate *k*_inact_. The K60P warhead is positioned much closer to the salt-bridge partners than that of XO44 and is more likely to affect, and be affected by, the dynamics of the associated structural components, particularly of the DFG-motif, suggesting that kinase activity state could be a major contributor to probe labeling. To explore the dynamics of the ligand and binding pocket prior to the formation of the reactive complex, and to ascertain any sensitivity to the protein conformation, we conducted a series of unbiased, 4 µs MD simulations of both K60P and XO44 probes reversibly associated but before covalent engagement (unlinked) with ABL1 and SRC kinases, including DFG-in and DFG-out conformations. The results of the simulations are summarized in Figure 7, as Empirical Cumulative Distribution Function (ECDF) plots (Figure S12) and discussed further in the Supplementary Molecular Modeling Discussion. Across these simulations, we observed increased reaction-ready proximity (warhead-S to lysine-N distances of < 4.5 Å) in the DFG-in conformations for both K60P and XO44, relative to the DFG-out conformations (Figure 7A). This preference can be explained by the observation that the DFG-out phenylalanine largely blocks proximity between the lysine and warhead of both K60P and XO44 when bound to the DFG-out conformation of ABL1 (Figure 7B), suggesting that each probe is more likely to label kinases in an ‘active’ conformation. Comparing the dynamics of ABL1 and SRC in the DFG-in conformation revealed that, while the K60P warhead maintains overall proximity to the catalytic lysine, it exhibits a reduced probability of achieving a reaction-ready proximity relative to XO44. This appears to be because of over-confinement of the warhead within the narrow kinase cleft, which restricts lysine rotameric flexibility and the approach geometry required for nucleophilic attack (Figure 7C).

**Figure 7:**
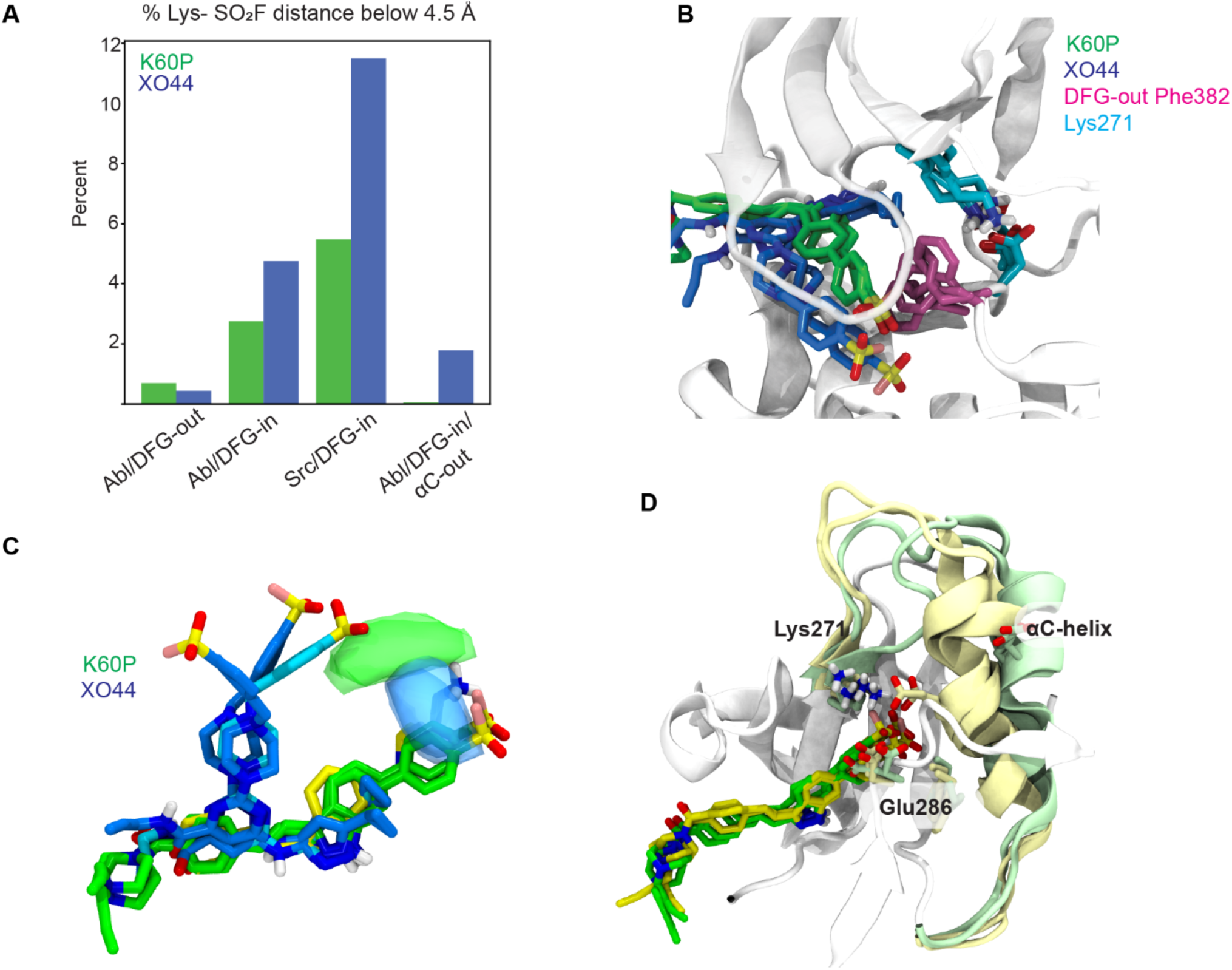
Molecular dynamics reveals preference of both K60P and X4K for DFG-in active conformation of kinases. (**A**) Estimation of the reaction-ready population in each system, determined by percentage of frames with < 4.5 Å Lysine to Warhead-S distance in each 4 µs simulation, excluding the first 100 ns. (**B**) Final structures of K60P (green) and XO44 (blue) in Abl/DFG-out simulations (both replicates overlayed). The DFG-out phenylalanine (pink) fences the warhead of both ligands away from the lysine (cyan, salt-bridging with DFG-Asp) and displaces the K60P core. (**C**) Positions of probes and the differing regions of highest lysine occupancy (blobs) for K60P (green) and XO44 (blue) for each of 2 replicates of ABL/DFG-in simulations at ∼4 µs. Expected poses for KW-2449 and lysine (yellow), and XO44 (cyan) are shown for reference. (**D**) Breaking of the Lys-Glu salt bridge and αC-helix adopting the out conformation in K60P ABL/DFG-in simulations. Two replicates are shown at both 250 ns (yellow) and at 500 ns (green).

The dynamics also inform us about the likelihood of a neutral reaction-ready lysine through the dependence of its p*K*_a_ on the protein conformation. Increased sampling of the DFG-out conformation, which may have a lower Lysine p*K*_a_ with ligand bound, is a plausible mechanism to increase the likelihood of reaction-ready lysine^40^. Both ligands showed a preference for the DFG-in conformation, but XO44 has been co-crystallized with other kinases in the DFG-out conformation^14^, and in ABL1 the extended P-loop conformation that it binds to is associated with the inactive state^41^. The K60P complex may also increase the likelihood of reaction-ready lysine by another mechanism, as simulations of the DFG-in conformation showed it could break the Lys to αC-helix-Glu salt-bridge by inducing the αC-helix-out conformation (Figure 7D), and also to weaken through sterics the salt-bridge between the DFG-Asp and the lysine. Other chemical steps may also be relevant^42^, a second deprotonation step of the intermediate complex also has potential to be rate-limiting and may be more easily deprotonated for XO44 where the larger unlinked warhead to lysine distance results in the reaction intermediate being further away from the salt-bridge partners. There are multiple possible reaction pathways, which could be investigated via QM/MM simulations^43^.

In summary, the overall slower labeling observed for K60P compared to X4K is likely a combination of both unexpected but lower intrinsic reactivity, and lower sampling of reaction-ready warhead-lysine proximities. But these differences are relatively small compared to the differences in modeled *k*_inact_. It is likely that the exceptional kinetic performance of X4K (notably over very similar analogs^31^) is due to subtle differences in the kinetics of nucleophilic attack by Lys271 and the subsequent steps of deprotonation and fluoride expulsion. These findings also align with the notion that kinetically tuned electrophilic probes can show higher on-target labeling that is resistant to quenching by high abundance off-target proteins in whole proteome or live cells. Overall, the apparent preference for DFG-in kinase conformations by both probes supports their utility as activity probes in cellular settings and warrants more global investigation across diverse kinases.

## Discussion

Here, we aimed to apply docking and molecular dynamics simulations to identify new covalent kinase activity probe scaffolds and to understand the contributors to probe specificity and kinetic efficiency. We focused on the hinge-binding indazole pharmacophore found in the clinical candidates KW-2449 and axitinib, under the premise that these molecules have a relatively promiscuous *in vitro* kinase inhibition profile and thus could serve as general kinome-targeted activity probes in lysates, live cells and tissues. Through a computational docking approach, we identified a subset of phenylsulfonylfluoride containing analogs for further study, including a cell-active probe, K60P. Direct target-engagement and competition profiling proteomics demonstrated that K60P and its progenitor KW-2449 broadly target kinases as well as targets in related nucleotide-binding pathways. These targets and profiles could be missed by focused recombinant profiling assays. This point is further emphasized by the fact that KW-2449 only weakly inhibited native ABL1 in cells (1.5-fold competed), despite being stated as a clinical development target. This discrepancy can be attributed to the competition with intracellular ATP in live cells as well as the difference between recombinant catalytic domains and full-length, native proteins in cellular environments^44^. These data add weight to the general consensus supporting the implementation of probe selectivity profiling beyond biochemical assays throughout the therapeutic discovery pipeline^45, 46^.

Cellular profiling and computational modeling studies demonstrated that K60P engages more than 100 kinases across three model cell lines and though it is not as promiscuous as XO44, K60P expands coverage across several kinase sub-families. Together, these probes expand the scope of targeted kinases by cell-permeable kinase probes (Figure 2). Our biochemical, proteomics and kinetic modeling studies make a clear case that the rapid X4K labeling of ABL1 relative to K60P is due to a faster *k*_inact_, and not a reversible binding affinity step. This result falls in line with recent studies that suggest that, beyond suitable non-covalent affinity and warhead reactivity, optimization of covalent bond formation through increasing the residence time of a ligand ‘reaction ready’ conformation is a primary contributor to potent and specific target labeling by covalent probes^35, 47, 48^. Here, we extend this to suggest that increased kinetic efficiency of covalent labeling increases the utility of kinase probes across diverse kinases, as seen here between K60P (lower kinetic efficiency, lower global kinome labeling) and XO44 (high kinetic efficiency, increased global kinome labeling). The increased performance of kinetically efficient probes likely involves less competition and absorption by abundant off-target proteins sites in the proteome, resulting in increased ‘on-target’ protein labeling within the kinome. Additionally, molecular dynamics simulations demonstrated that K60P and XO44 (or its biotin probe, X4K) prefer the active, DFG-in conformations of ABL1 and SRC, which may be more general in the kinome. These data supports their utility for comparative kinase profiling in cell and patient samples, inhibitor selectivity profiling, as well as more focused spatial and single-cell activity profiling studies^16–19^. Finally, we posit that the combination of computational docking and molecular dynamics simulation described here can be used more generally to identify suitable ligand scaffolds for synthesis and testing, followed by optimization by kinetic modeling to identify more efficient paths to reaction-ready conformations.

## Code Availability

A Jupyter notebook for kinetic modeling is accessible at github repository: https://github.com/RouxLab/covalent-kinase-activity-probes-2025

## Supporting Information

- All experimental details, materials and methods, ^1^H-NMR spectra, supplementary molecular modeling methodology, supplementary molecular modeling discussion, supplementary kinetic modeling methodology, supplementary kinetic modeling discussion, supplementary figures and supplementary Table S4
- Table S1-S3: Raw proteomics data for K60P-Bio, X4K, and KW-2449 competition with K60P-Bio

## Acknowledgements

We thank S. Ahmadiantehrani for her valuable suggestions during the writing process, figure drafting, and proofreading assistance.

## Author Information

**Corresponding Authors**

**Raymond E. Moellering -** Department of Chemistry and Institute for Genomics and Systems Biology, The University of Chicago, 929 East 57th Street, Chicago, Illinois 60637, United States;

**Benoît Roux -** Department of Chemistry and Department of Biochemistry and Molecular Biophysics, The University of Chicago, 929 East 57th Street, Chicago, Illinois 60637, United States;

**Authors**

**Pratyasha Chakraborty** - Department of Chemistry, The University of Chicago, 929 East 57th Street, Chicago, Illinois 60637, United States

**Anthony Carlos** - Department of Chemistry, The University of Chicago, 929 East 57th Street, Chicago, Illinois 60637, United States

**Trayder Thomas** - Department of Biochemistry and Molecular Biophysics, The University of Chicago, 929 East 57th Street, Chicago, Illinois 60637, United States

**Kyle Ghaby** - Department of Biochemistry and Molecular Biophysics, The University of Chicago, 929 East 57th Street, Chicago, Illinois 60637, United States

**Shaghayegh Fathi** - Department of Chemistry, The University of Chicago, 929 East 57th Street, Chicago, Illinois 60637, United States

**Mukta Sharma** - Department of Biochemistry and Molecular Biophysics, The University of Chicago, 929 East 57th Street, Chicago, Illinois 60637, United States

**Ngoc Kim Nguyen** - Department of Chemistry, The University of Chicago, 929 East 57th Street, Chicago, Illinois 60637, United States

**Mason Farmwald** - Department of Chemistry, The University of Chicago, 929 East 57th Street, Chicago, Illinois 60637, United States

**Lydia Blachowicz** - Department of Biochemistry and Molecular Biophysics, The University of Chicago, 929 East 57th Street, Chicago, Illinois 60637, United States

**Emma Alcyone** - Department of Chemistry, The University of Chicago, 929 East 57th Street, Chicago, Illinois 60637, United States

**Shaopeng Yu** - Department of Chemistry, The University of Chicago, 929 East 57th Street, Chicago, Illinois 60637, United States

**Kavya Smitha Pillai** - Department of Chemistry, The University of Chicago, 929 East 57th Street, Chicago, Illinois 60637, United States

## Author contributions

R.E.M. and B.R. conceptualized the study. P.C., A.C., and S.F. synthesized the probes. T.T. and K.G. performed molecular dynamics simulations. T.T. and M.S. performed docking studies. P.C. carried out mass spectrometry experiments. P.C. and A.C. performed bioinformatics analysis for proteomics experiments. K.G. and T.T. performed kinetic modeling. P.C., N.K.N., M.F. performed biochemical assays. L.B. performed protein purification. E.A., K.S.P., S.Y. helped with the syntheses. P.C., A.C., and T.T. contributed equally to this work. P.C., A.C., T.T., K.G., R.E.M., and B.R. wrote the manuscript. R.E.M. and B.R. are co-corresponding authors. All authors reviewed the manuscript.

