## Supplementary Information for "Molecular Dynamics-Guided Design and Chemoproteomic Profiling of Covalent Kinase Activity Probes"

### Supporting Information for Molecular Dynamics-Guided Design and Chemoproteomic Profiling of Covalent Kinase Activity Probes

**Pratyasha Chakraborty[†, 1], Anthony Carlos[†, 1], Trayder Thomas[†, 2], Kyle Ghaby[2], Shaghayegh Fathi[1], Mukta Sharma[2], Ngoc Kim Nguyen[1], Mason Farmwald[1], Lydia Blachowicz[2], Shaopeng Yu[1], Kavya Smitha Pillai[1], Benoît Roux[*, 1,2], Raymond Moellering[*,1,3]**

**[1]Department of Chemistry, [2]Department of Biochemistry and Molecular Biophysics, & [3]Institute for Genomics and Systems Biology, The University of Chicago. Chicago, IL, 60637, USA.**

[†] These authors contributed equally.

#### Table of Contents

Supplementary Figures and Tables

Methods for Biology

Supplementary Molecular Modeling Methodology

Supplementary Molecular Modeling Discussion

Supplementary Kinetic Modeling Methodology

Supplementary Kinetic Modeling Discussion

General Information for Chemistry

Preparation of probes

$^1$H NMR Spectra of probes

References

### Supplementary Figures and Tables:

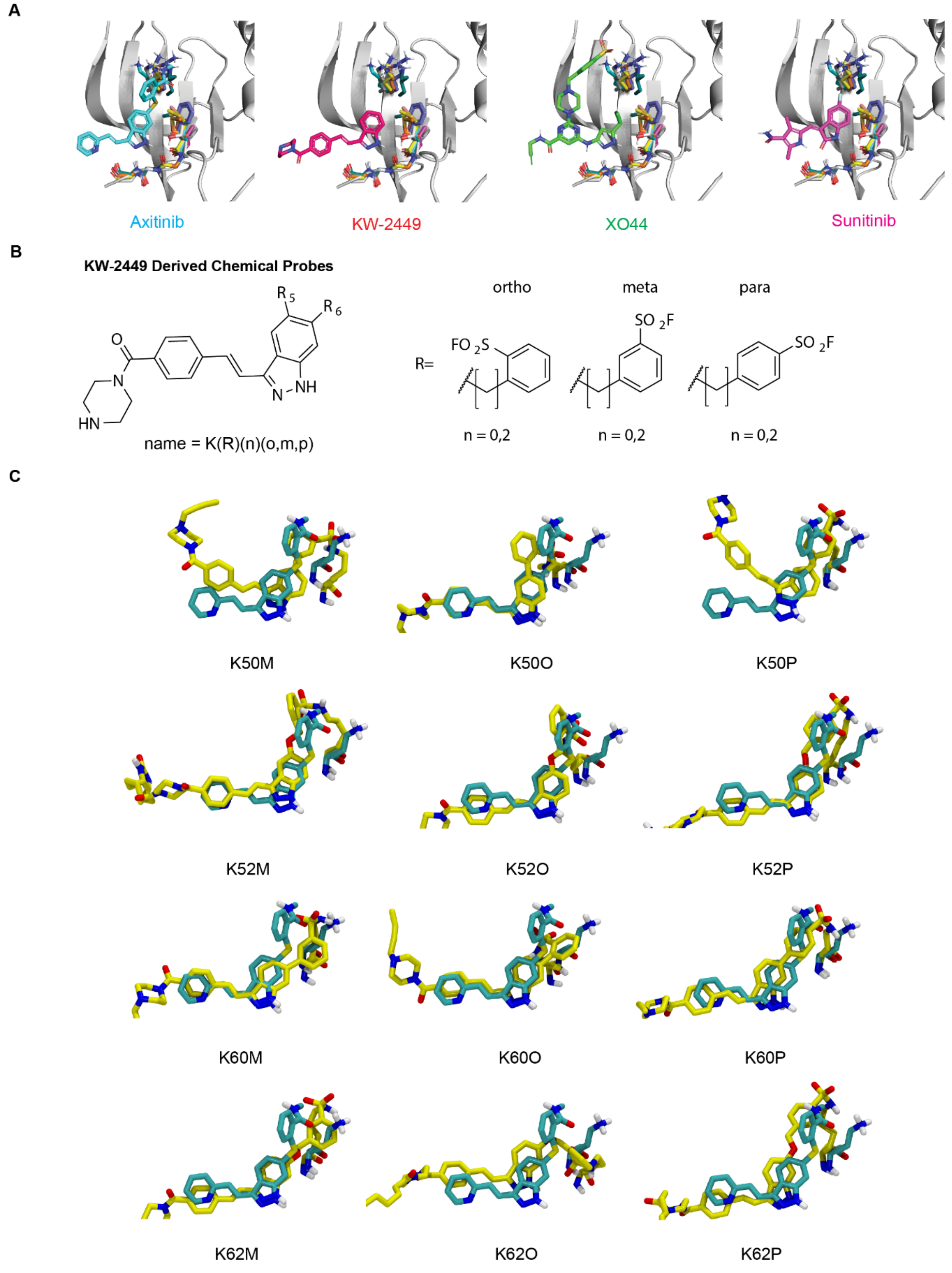

**Figure S1: Modeled poses of starting scaffolds and final linked poses of screened compounds** (**A**) Overlayed binding poses of relevant ligands (shown separately) alongside overlayed conserved binding pocket features from each of 7 kinase families (**B**) Naming convention for KW-2449 scaffold probes in molecular dynamics screening. (**C**) Final structures of KW-2449 scaffold probes and lysine in MD screening simulations (yellow) after 100 ns, overlayed with co-crystallized poses of axitinib and lysine from PDB ID: 4WA9 (cyan).

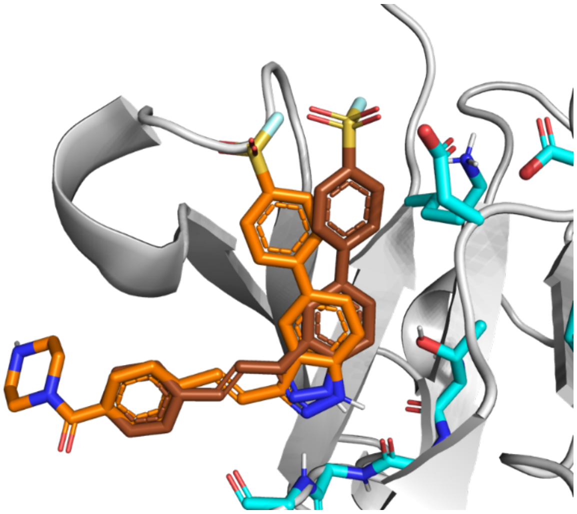

**Figure S2: Alternative alkene rotation for K5* series compounds**. Modeled pose of K50P (orange) aligned to axitinib in PDB ID: 4TWP, and alternative rotamer of the ring-connecting alkene (brown). The backbone of kinase hinge residues, gatekeeper residue, conserved lysine, and its salt-bridge partners are shown in cyan.

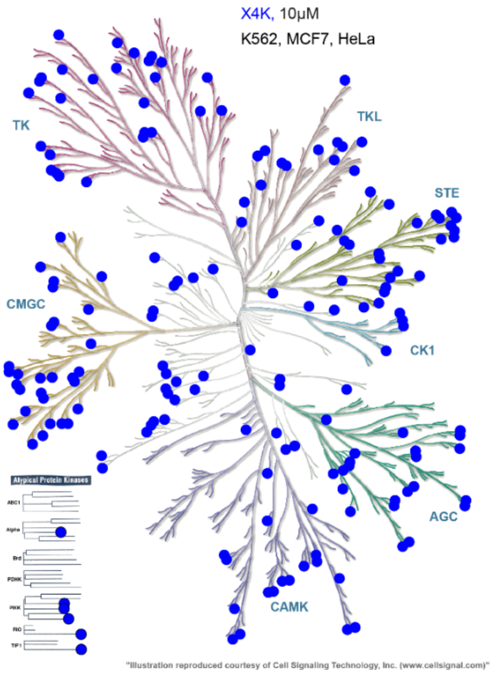

**Figure S3: Kinome scope of X4K**. Dendogram representation of the kinases targeted by X4K in K562, Hela, and MCF7 cell lines shown as blue spheres.

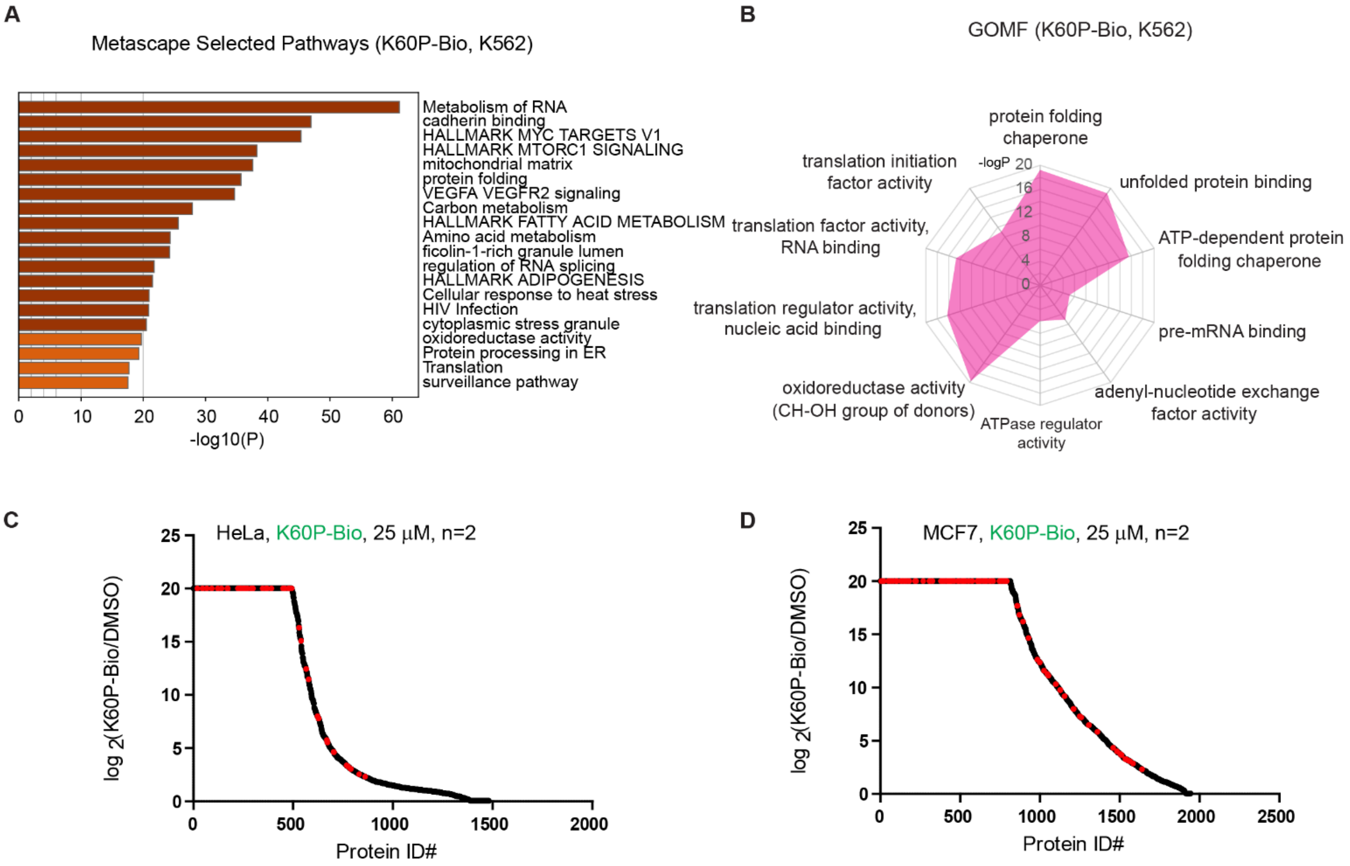

**Figure S4: K60P-Bio labels kinases and other classes of proteins across different cell lines.** (**A**) Metascape analysis of proteins enriched more than 2-fold by K60P-Bio in K562 cells. (**B**) Gene Ontology Molecular Function analysis of proteins enriched more than 2-fold by K60P-Bio in K562 cells. (**C-D**) Waterfall plot of proteins enriched by K60P-Bio (25µM, 4 hours) in live HeLa (C) and MCF7 (D) cells. Kinases are shown as red dots. Control cells were treated with DMSO. Cutoff: SILAC ratio >2; n=2

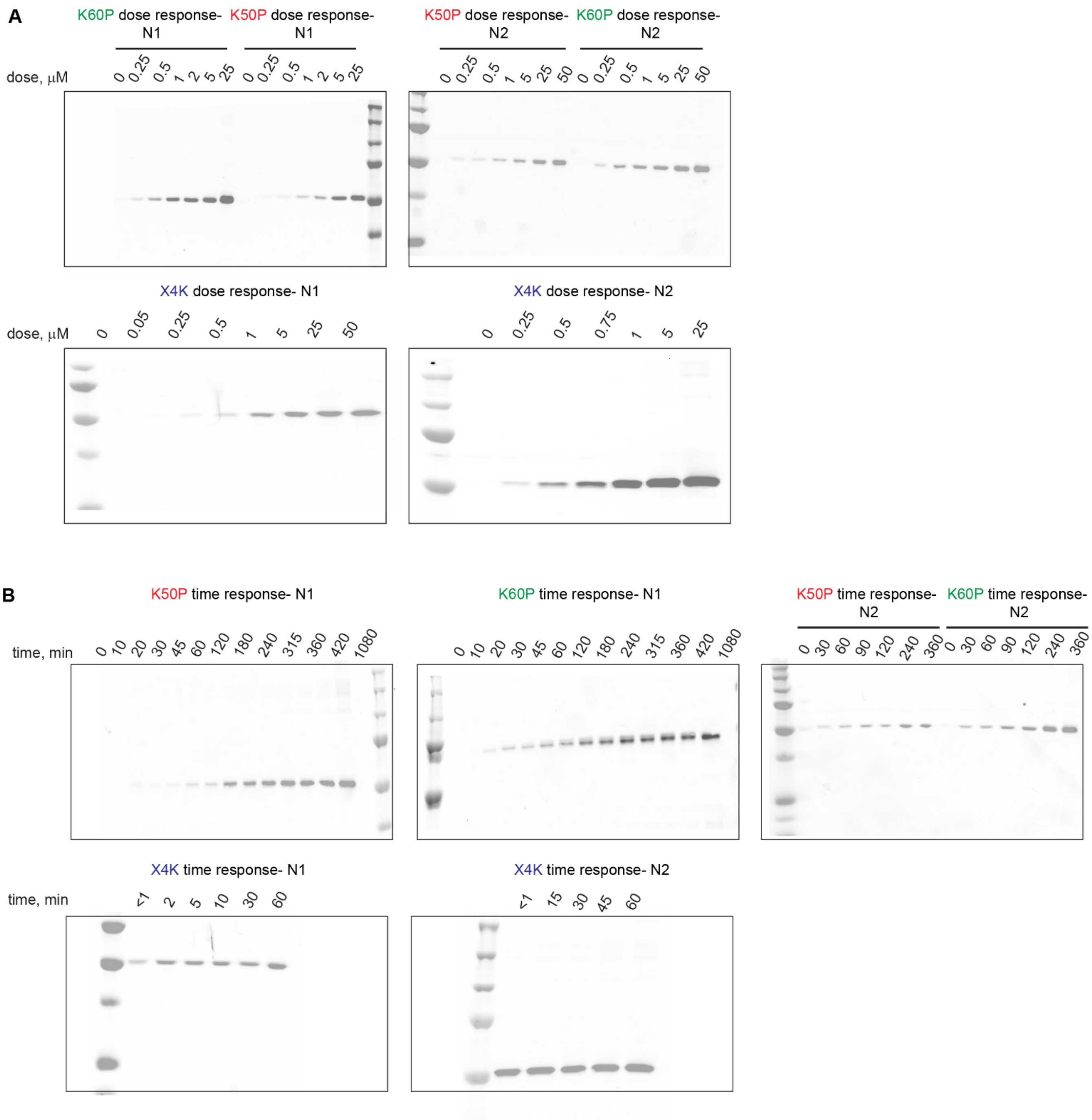

**Figure S5: Uncropped blots for dose- and time- responses of ABL1 labeling by K60P, K50P, and X4K**- bioreplicates of Figure 4A and 4C- dose response (**A**) and time response (**B**) of ABL1 labeling by K60P, K50P, and X4K

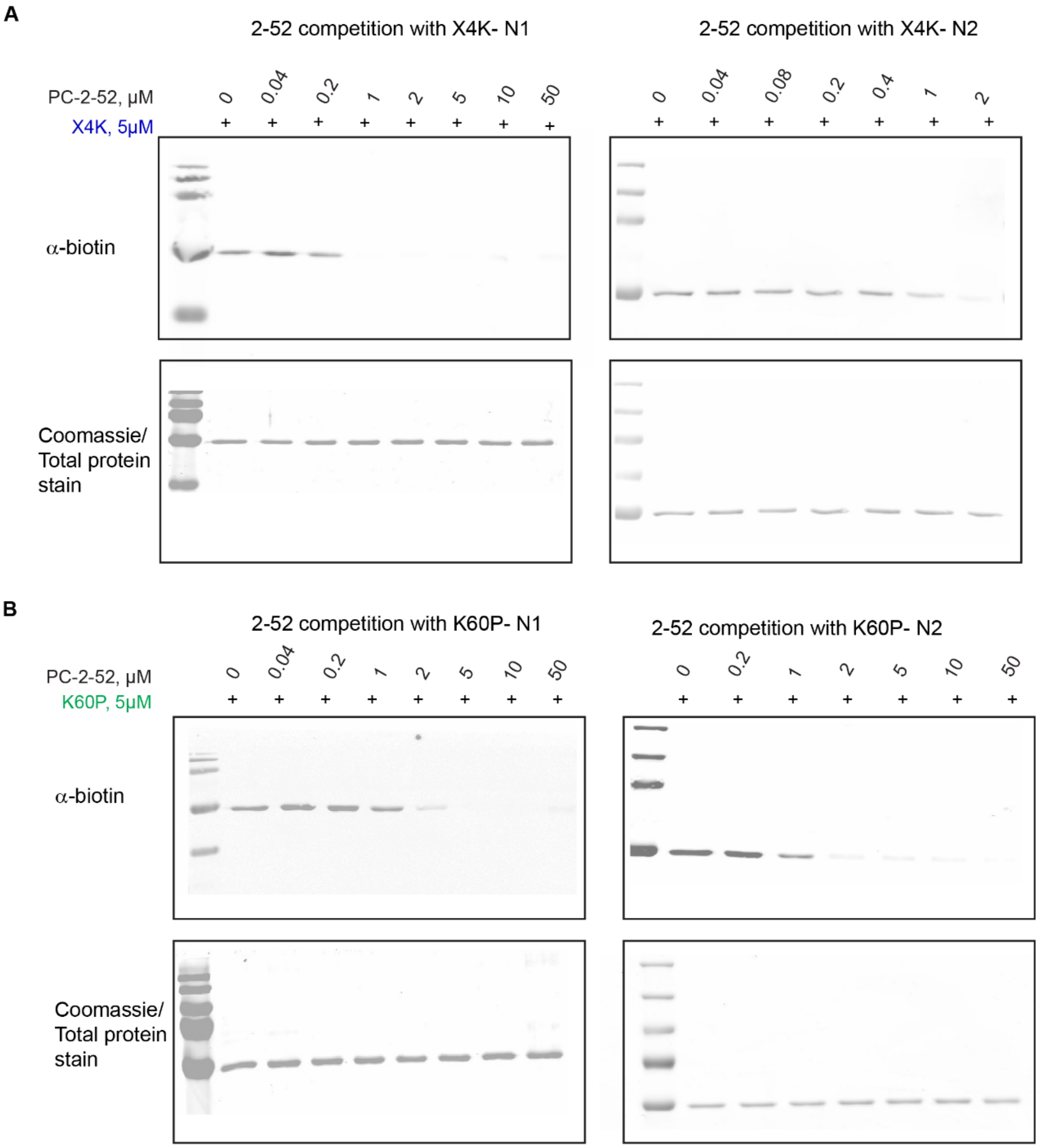

**Figure S6: Uncropped blots for competition of PC-2-52 with X4K and K60P-** bioreplicates of Figure 4B- competition of PC-2-52 with X4K (**A**) and with K60P (**B**) for ABL1 labeling

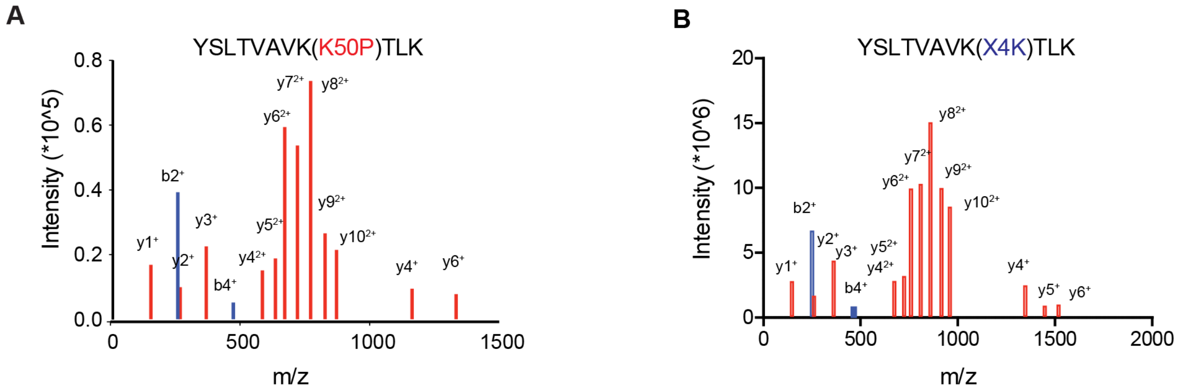

**Figure S7A: K50P and X4K covalently label the conserved K271 of ABL1**. MS/MS spectra of K271 labeled peptides for K50P (**A**) and X4K (**B**).

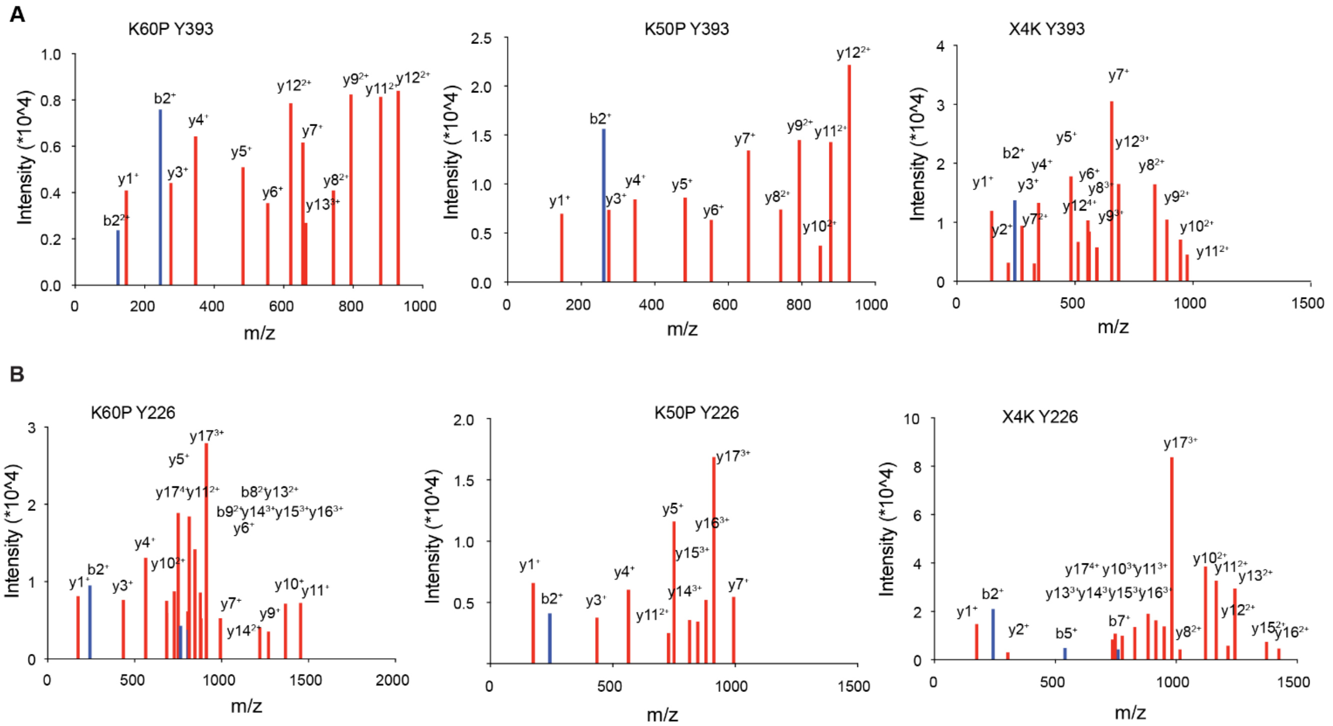

**Figure S7B: K60P, K50P, and X4K covalently label Y393 and Y226 in ABL1**. MS/MS spectra of Y393 (**A**) and Y226 (**B**) labeled peptides for K60P (left), K50P (middle), and X4K (right).

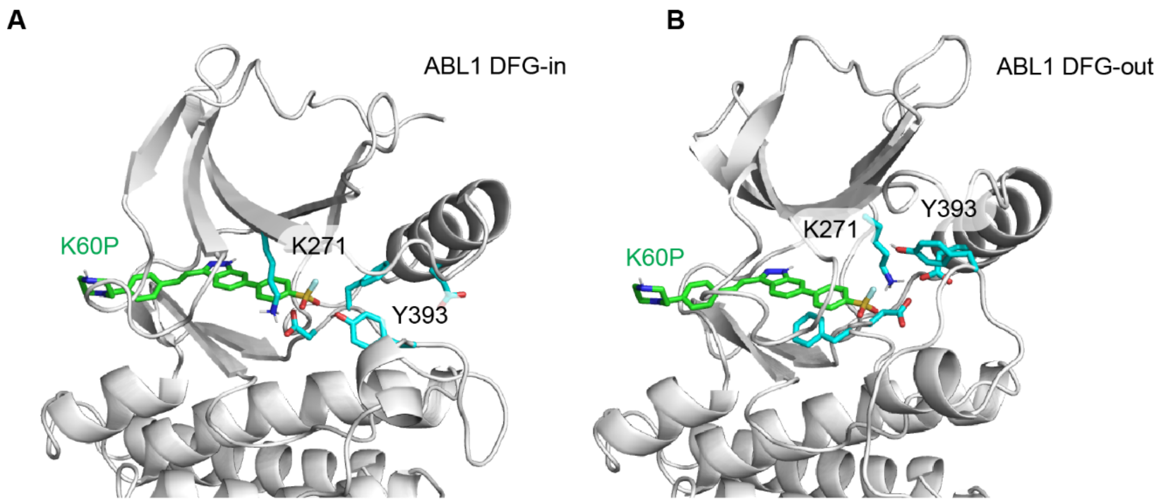

**Figure S7C: Conformations of apo-ABL1 identified with Y393 closest to K60P.** K60P is modeled in green with DFG-in (**A**) and DFG-out (**B**) conformations. K271 and Y393 are shown in cyan.

| | K50P | | | | K60P | | XO44 | |
|---|---|---|---|---|---|---|---|---|
| | $Mod_{del}$ | | $Mod_{sil}$ | | | | | |
| $k_{on}$ $(\frac{1}{\mu MS})$ | $1.82 \times 10^{-2}$ | * | $1.82 \times 10^{-2}$ | * | $1.82 \times 10^{-2}$ | * | $1.26 \times 10^{-3}$ | $5.20\times10^{-3}$<br>$4.56\times10^{-4}$ |
| $k_{off}$ $(\frac{1}{S})$ | $7.62 \times 10^{-2}$ | $1.50\times10^{-1}$<br>$2.29\times10^{-2}$ | $1.17 \times 10^{-1}$ | $2.25\times10^{-1}$<br>$5.20\times10^{-2}$ | $2.10 \times 10^{-2}$ | $4.54\times10^{-2}$<br>$5.80\times10^{-3}$ | $8.95 \times 10^{-2}$ | * |
| $K_d$ (μM) | $4.19 \times 10^{0}$ | $8.27\times10^{0}$<br>$1.26\times10^{0}$ | $6.43 \times 10^{0}$ | $1.24\times10^{1}$<br>$2.86\times10^{0}$ | $1.15 \times 10^{0}$ | $2.50\times10^{0}$<br>$3.19\times10^{-1}$ | $7.13 \times 10^{1}$ | $1.51\times10^{2}$<br>$7.89\times10^{0}$ |
| $k_{inact}$ $(\frac{1}{S})$ | $1.19 \times 10^{-4}$ | $1.64\times10^{-4}$<br>$7.40\times10^{-5}$ | $2.06 \times 10^{-4}$ | $2.82\times10^{-4}$<br>$1.41\times10^{-4}$ | $1.91 \times 10^{-4}$ | $2.45\times10^{-4}$<br>$1.40\times10^{-4}$ | $1.34 \times 10^{-1}$ | $3.12\times10^{-1}$<br>$5.99\times10^{-3}$ |
| $K_I$ (μM) | $4.19 \times 10^{0}$ | $8.28\times10^{0}$<br>$1.26\times10^{0}$ | $6.44 \times 10^{0}$ | $1.24\times10^{1}$<br>$2.87\times10^{0}$ | $1.16 \times 10^{0}$ | $2.51\times10^{0}$<br>$3.27\times10^{-1}$ | $1.86 \times 10^{2}$ | $5.85\times10^{2}$<br>$8.45\times10^{0}$ |
| $C_{eff}$ $(\frac{1}{\mu MS})$ | $2.82 \times 10^{-5}$ | $5.03\times10^{-5}$<br>$1.55\times10^{-5}$ | $3.18 \times 10^{-5}$ | $5.32\times10^{-5}$<br>$1.70\times10^{-5}$ | $1.63 \times 10^{-4}$ | $2.92\times10^{-4}$<br>$6.85\times10^{-5}$ | $6.43 \times 10^{-4}$ | $1.35\times10^{-3}$<br>$3.32\times10^{-4}$ |
| $k_{ns}$ $(\frac{1}{\mu MS})$ | $5.69 \times 10^{-7}$ | $1.43\times10^{-6}$<br>$1.46\times10^{-9}$ | $5.38 \times 10^{-7}$ | $1.53\times10^{-6}$<br>$1.85\times10^{-9}$ | $1.65 \times 10^{-6}$ | $2.22\times10^{-6}$<br>$1.11\times10^{-6}$ | $5.14 \times 10^{-6}$ | $6.86\times10^{-6}$<br>$3.36\times10^{-6}$ |

| | | | | | | | |
|---|---|---|---|---|---|---|---|
| $k_{\mathrm{del}}\ (\frac{1}{s})$ | $1.12 \times 10^{-4}$ | $1.71\times10^{-4}$<br>$5.44\times10^{-5}$ | | | | | |
| $\mathrm{Frac_{sil}}$ | | | $2.79 \times 10^{-1}$ | $4.21\times10^{-1}$<br>$1.28\times10^{-1}$ | | | |
| $\lambda_{\mathrm{tr,BR1}}$ | $1.56 \times 10^{0}$ | | $1.46 \times 10^{0}$ | | $6.65 \times 10^{-1}$ | | $5.93 \times 10^{-1}$ |
| $\lambda_{\mathrm{tr,BR2}}$ | $1.37 \times 10^{0}$ | | $1.28 \times 10^{0}$ | | $8.01 \times 10^{-1}$ | | |
| $\lambda_{\mathrm{dr,BR3}}$ | $2.40 \times 10^{0}$ | | $2.34 \times 10^{0}$ | | $1.29 \times 10^{0}$ | | $5.33 \times 10^{-1}$ |
| $\lambda_{\mathrm{dr,BR4}}$ | $3.05 \times 10^{0}$ | | $2.92 \times 10^{0}$ | | $1.50 \times 10^{0}$ | | |

**Table S4. The estimated median and 95% highest density interval ($\left(\begin{smallmatrix}\text{upper}\\ \text{lower}\end{smallmatrix}\right)$) of parameters from BMMC simulations.** Parameters without a range were fixed during BMMC. Values marked with an asterisk (*) were also fixed during LS minimization. The $K_d$, $K_I$, and $C_{eff}$ were not explicit parameters, and instead calculated per sample.

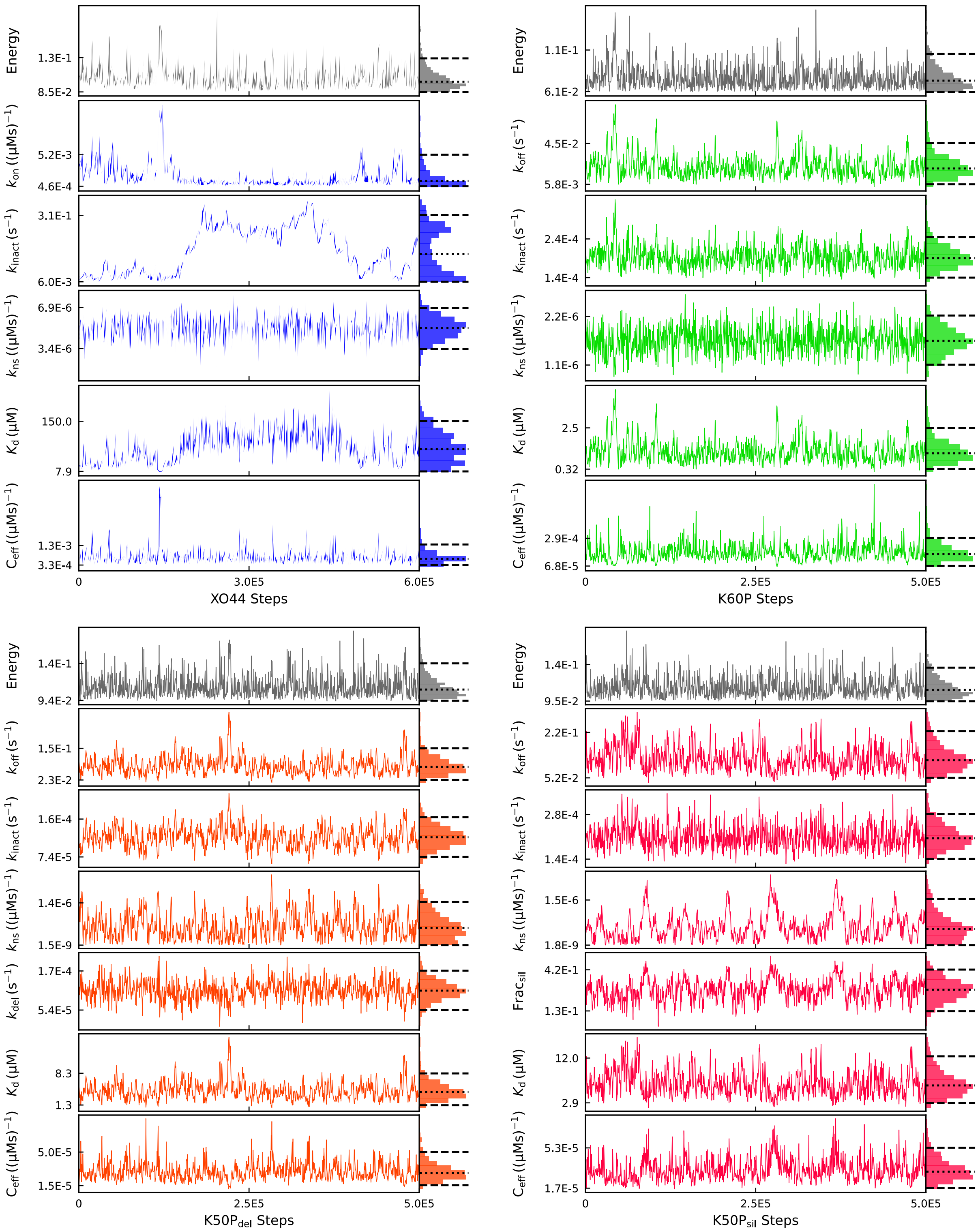
Energy
1.3E-1
8.5E-2
$k_{on}$ ((μMs)$^{-1}$)
5.2E-3
4.6E-4
$k_{inact}$ (s$^{-1}$)
3.1E-1
6.0E-3
$k_{ns}$ ((μMs)$^{-1}$)
6.9E-6
3.4E-6
$K_d$ (μM)
150.0
7.9
$C_{eff}$ ((μMs)$^{-1}$)
1.3E-3
3.3E-4
0
3.0E5
6.0E5
XO44 Steps
Energy
1.1E-1
6.1E-2
$k_{off}$ (s$^{-1}$)
4.5E-2
5.8E-3
$k_{inact}$ (s$^{-1}$)
2.4E-4
1.4E-4
$k_{ns}$ ((μMs)$^{-1}$)
2.2E-6
1.1E-6
$K_d$ (μM)
2.5
0.32
$C_{eff}$ ((μMs)$^{-1}$)
2.9E-4
6.8E-5
0
2.5E5
5.0E5
K60P Steps
Energy
1.4E-1
9.4E-2
$k_{off}$ (s$^{-1}$)
1.5E-1
2.3E-2
$k_{inact}$ (s$^{-1}$)
1.6E-4
7.4E-5
$k_{ns}$ ((μMs)$^{-1}$)
1.4E-6
1.5E-9
$k_{del}$ (s$^{-1}$)
1.7E-4
5.4E-5
$K_d$ (μM)
8.3
1.3
$C_{eff}$ ((μMs)$^{-1}$)
5.0E-5
1.5E-5
0
2.5E5
5.0E5
K50P$_{del}$ Steps
Energy
1.4E-1
9.5E-2
$k_{off}$ (s$^{-1}$)
2.2E-1
5.2E-2
$k_{inact}$ (s$^{-1}$)
2.8E-4
1.4E-4
$k_{ns}$ ((μMs)$^{-1}$)
1.5E-6
1.8E-9
Frac$_{sil}$
4.2E-1
1.3E-1
$K_d$ (μM)
12.0
2.9
$C_{eff}$ ((μMs)$^{-1}$)
5.3E-5
1.7E-5
0
2.5E5
5.0E5
K50P$_{sil}$ Steps

**Figure S8: Time-evolution of the effective energy and variable parameters during the BMMC simulation.** Samples were collected every 500 steps to account for autocorrelation of parameters. The equilibrium non-covalent dissociation constant ($K_d = k_{\text{off}}/k_{\text{on}}$) and covalent efficiency ($C_{\text{eff}} = k_{\text{inact}}/K_{\text{I}}$, where $K_{\text{I}} = (k_{\text{off}} + k_{\text{inact}})/k_{\text{on}}$) were calculated at each sample for K60P (green), K50P (orange, red), and X4K (blue).The distributions of each time evolution are shown to the right of each plot. The median of each parameter is marked with a dotted line, and the bounds of the 95% highest density interval are marked with a dashed line.

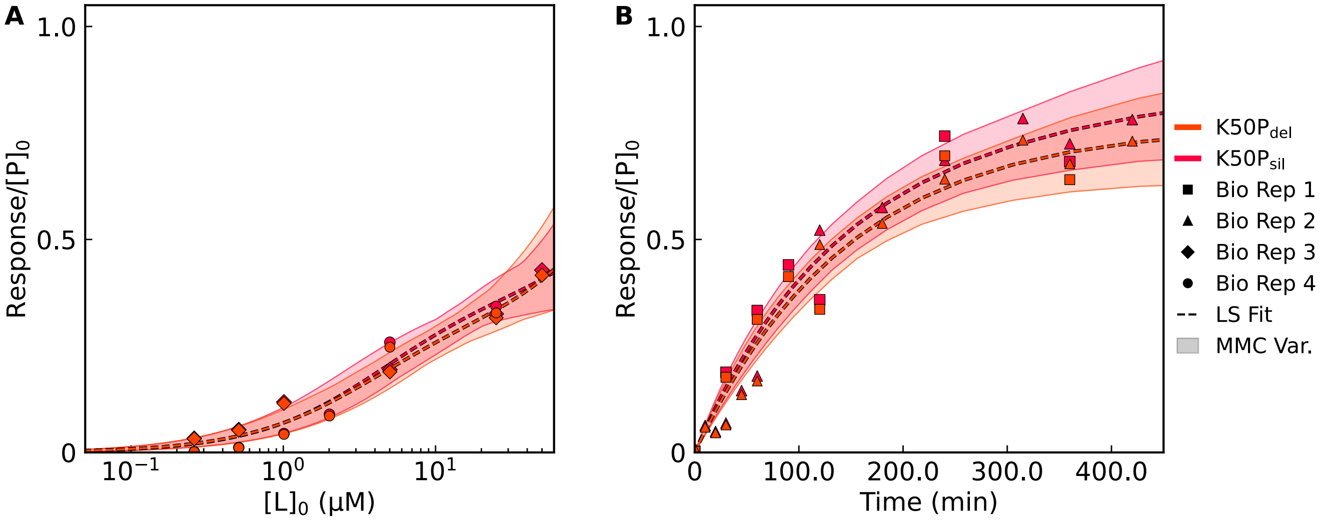

**Figure S9: The least-squares (LS) fit and Metropolis Monte-Carlo (BMMC) variation of dose-responses (A) and time-responses (B) for each K50P modified scheme.** The samples have been filtered so that each kinetic parameter falls within its respective 95% highest density interval. Simulated "response" is the sum of the specifically- and nonspecifically-bound covalent complexes.

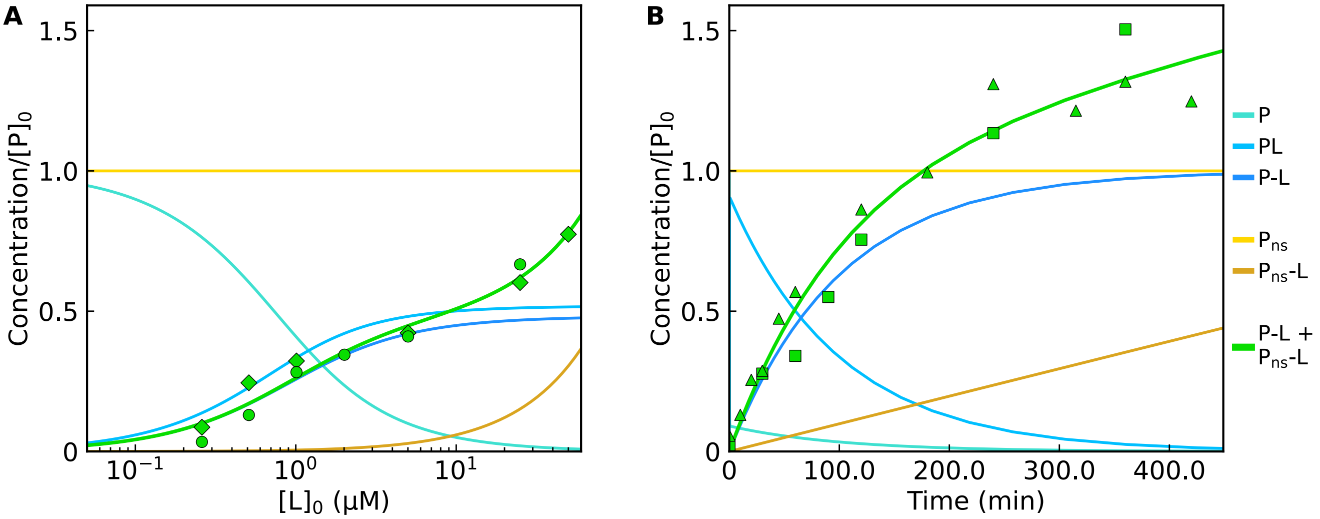

**Figure S10: Evolution of species over time during a simulated dose-response (A) and time-response (B) using the best parameter values from the least-squares fit for K60P.** The normalized concentrations of all Scheme 1 species, the normalized sum of both covalently bound species, and the normalized experimental data are shown.

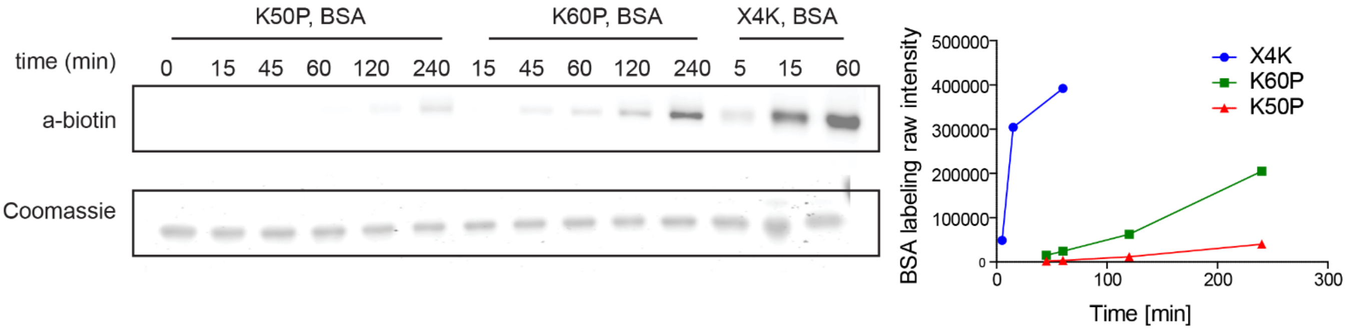

**Figure S11: X4K, K60P, and K50P exhibited different inherent reactivities.** Rate of non-specific target protein BSA with K60P, K50P, and X4K shows intrinsic higher reactivity for X4K compared to K50P and K60P

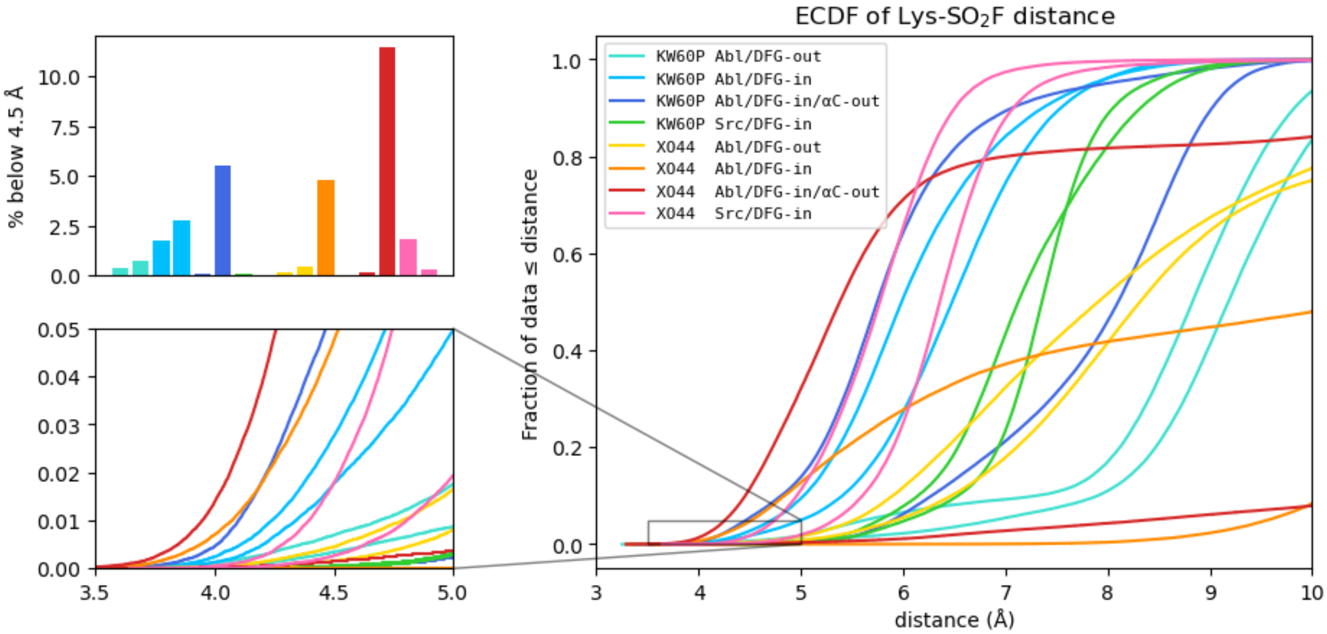

**Figure S12: Empirical Cumulative Distribution Function (ECDF) of the Lysine to Warhead-S distance in each 4 µs simulation, excluding the first 100 ns**. Duplicates with slight tweaks to starting conformation are shown as the same color.

#### Methods for Biology

##### Cell culture

K562, HeLa, and MCF7 cells were propagated in RPMI (Corning) supplemented with 10% fetal bovine serum (FBS, Corning) and 1% penicillin/streptomycin (Gibco). All cell lines were grown at 37℃ in a 5% CO2 humidified incubator. All cell lines tested negative for mycoplasma using Lonza MycoAlert PLUS Detection Kit.

##### In-vitro recombinant Abl labeling assays

The c-Abl three-domain construct (SH3-SH2-kinase; residues 46-515, human Abl 1a numbering) was coexpressed in E. coli BL21(DE3) cells with YopH phosphatase and purified as described in Seeliger et al.[1] with the following modifications. Cleared cell lysate in 50 mM Tris (pH 8.0), 500 mM NaCl, 5% glycerol, 25 mM imidazole (buffer A) was loaded onto Ni-NTA agarose resin (Qiagen) equilibrated with buffer A. The resin was washed with 15 bed volumes of buffer A and eluted with 3 bed volumes of buffer B (buffer A plus 0.5 M imidazole). The His tag was cleaved from the eluted kinase by incubation with 1 mg of TEV protease per 25 mg of kinase while dialyzing against 20 volumes of 20 mM Tris (pH 8.0), 100 mM NaCl, 5% glycerol, 1 mM DTT for 16 h at 4°C with a 13 kDa molecular weight cutoff (MWCO) membrane. In order to separate the kinase from the cleaved His tag, TEV protease (which contains a His tag) and YopH phosphatase (which binds nonspecifically to the Ni-NTA resin), the dialysate was passed over Ni-NTA agarose resin equilibrated with buffer A. The flow-through contained pure kinase protein with the His tag removed. Concentration of the purified kinase was determined by absorbance spectroscopy at 280 nm using the calculated extinction coefficient of 60,550 $M^{-1}$ $cm^{-1}$.

The purified protein was buffer exchanged to PBS using 30kDa MWCO Millipore Amicon centrifugal filter (3000xg, 10min, 4°C, 8 times) to remove tris in the protein storage buffer from interacting with probe. The resulting protein concentration was measured using BCA assay and adjusted to make 0.025 mg/mL protein stock.

For dose response, 20 µL of protein stock was added to each sample followed by addition of 5 µL of probe stock (5X). The samples were incubated for 1 hour at 4°C. The samples were then diluted with 4X Laemmli buffer containing 50 mM dithiothreitol (DTT) as reducing agent, heated to 95℃ for 5 minutes, and cooled to room temperature. The samples were then resolved on a 10% SDS-PAGE gel and transferred to nitrocellulose membranes using standard western blotting techniques. Membranes were blocked with 2% BSA in TBS containing 0.1% Tween-20 (TBST) and probed with streptavidin-800 (Licor IRDye® 800CW, Licor Odyssey CLx imager) to detect probe labeling. Coomassie staining was done as loading control for the gels. Blot intensities were quantified using ImageJ and normalized in Microsoft Excel.

For time response, 20 µL of protein stock was added to each sample followed by addition of 5 µL of probe stock (5X, 50 µM). At the end of each time point, the respective sample was diluted with 4X Laemmli buffer containing 50 mM dithiothreitol (DTT) as reducing agent and heated to 95℃ for 5 minutes. The samples were then then probed for streptavidin signal according to western blot protocol mentioned above. Coomassie staining was done as loading control for the gels.

**K50P and K60P probe dosing on live cells and lysates for immunoblot analysis**

Wildtype HeLa cells were seeded in 24-well plate at 0.1*10^6 cells per well in 1mL of RPMI media (Corning) supplemented with 10% fetal bovine serum (Corning) and 1% penicillin/streptomycin (Gibco) 24 hour before experimentation. Cells were treated with 0-50 µM of probe in 500 µL of serum-free, phenol-red-free RPMI media (Gibco) for 4 hr at 37°C. The cells were then washed in 500uL PBS twice and lysed in ice-cold 1x RIPA buffer supplemented with protease inhibitor (Roche) and 1mM DTT. Cells were allowed to lyse while shaking on ice for 20 min. The lysates were collected and insoluble debris was cleared using centrifugation. The supernatant was diluted in 4X Laemmli buffer containing 50 mM dithiothreitol (DTT) as a reducing agent. SDS-PAGE samples were prepared by heating the mixture at 95 °C for 5 minutes, then cooling to room temperature. The samples were resolved on a 10% SDS-PAGE gel and transferred to nitrocellulose membranes using standard western blotting techniques. Membranes were blocked with 2% BSA in TBS containing 0.1% Tween-20 (TBST) and probed with streptavidin-800 (Licor IRDye® 800CW, Licor Odyssey CLx imager) to detect biotin/desthiobiotin labeling. Blot intensities were quantified using ImageJ and normalized in Microsoft Excel.

For lysate dosing studies, clarified HeLa lysates were treated with 0-50 µM probe for 2 hours at 37°C before preparing samples for immunoblotting as described above.

**ABL1 site of labeling assays**

For mapping the site of labeling of the probes in ABL1, the protein was buffer-exchanged to PBS as mentioned above and diluted to a concentration of 0.2 mg/mL protein in PBS. 250 µL of the protein was taken and treated with 10 µM probe for 1h at 37 °C. Excess probe was quenched with 10mM Tris base and 1X 8M urea (250 μL) was added. Then, DTT aqueous solution (5 μL, 1M) was added, followed by incubation at 65 °C for 15 min. Iodoacetamide aqueous solution (40 μL, 0.5M) was added to the sample, followed by further incubation for 30 min in the dark. The sample was then transferred to a 30kDa MWCO Millipore Amicon centrifugation filter unit and washed twice with 1mL 2M urea in 25mM ammonium bicarbonate solution. The sample volume was brought up to 300 μL with 2M urea in 25mM ammonium bicarbonate. $CaCl_2$ (3 μL, 100mM) was added to the sample followed by 1 μg of sequencing-grade trypsin. The samples were digested overnight at 37 ˚C while shaking. Following trypsinization, supernatant was collected, acidified with HPLC grade formic acid (2% final, pH 2-3), and peptides were then desalted on ZipTip C18 tips (100 μL, Millipore), dried under vacuum, resuspended with LC-MS grade water (Sigma Aldrich), and then lyophilized.

Lyophilized peptides were dissolved in LC-MS/MS Buffer ($H_2O$ with 0.1% formic acid, LC-MS grade, Sigma Aldrich) for proteomic analysis. Proteomic methods reported are adopted from our previous reports[2]. LC–MS/MS analysis for proteomics samples was performed with an UltiMate 3000 RSLCnano System (Thermo Fisher Scientific) using an Acclaim PepMap RSLC C18 column (75 μm × 15 cm, 2 μm, 100 Å, Thermo Fisher Scientific) with an in-line Acclaim PepMap 100 C18 trap column (75 μm × 2 cm, 3 μm, 100 Å, Thermo Fisher Scientific) heated to 45 °C. The LC system was coupled to an Orbitrap Exploris 480 and Nanospray Flex Ion Source with stainless steel emitter tip (Thermo Fisher Scientific). Mobile phase A was composed of $H_2O$ supplemented with 0.1% formic acid, and mobile phase B was composed of $CH_3CN$ supplemented with 0.1% formic acid. The instrument was run at 0.3 μl min−1 with 2-hour gradients. MS/MS spectra were collected for the entirety of the gradient using a data-dependent, 2-second cycle time setting with the following details: full MS scans were acquired at a resolution of 120,000, scan range of 380 m/z to 1,500 m/z, maximum IT of 25 ms, normalized AGC target of 300% and data collection in profile mode. MS2 scans were performed by high-energy collision dissociation (HCD) fragmentation with a resolution of 15,000, normalized AGC target of 50%, maximum IT of 50 ms, HCD collision energy of 30% and data collection in centroid mode. The isolation window for precursor ions was set to 1.6 m/z. Peptides with a charge state of 1, 7+ and unassigned were excluded, and dynamic exclusion was set to 40 seconds. The RF lens % was set to 40 with a spray voltage value of 2.0 kV and an ionization chamber temperature of 300 °C.

Data were processed using the SEQUEST HT search engine node within the Proteome Discoverer 3.0 software package. Data were searched using a concatenated target/decoy UniProt database of the human proteome with isoforms. Digest enzyme specificity was set to trypsin with up to two missed cleavages allowed, and peptide length was set to between 6 and 144 residues. Precursor mass range was set to 350–6500. Precursor mass tolerance was set to 10 ppm, and fragment mass tolerance was set to 0.02 Da. Up to 4 dynamic modifications were allowed per peptide, including K60P/K50P (+ 666.262) or X4K (+ 858.353) modification on Lysine or Tyrosine, oxidized methionine (+15.9949), N-terminal acetylation (+42.0106), N-terminal Met-loss (−131.0405) and N-terminal Met-loss + acetylation (−89.0299). Cysteine carboxyamidomethylation (+57.0215) was set as a static modification. A minimum of two peptides, with a minimum length of 6, was required for protein identification, and false discovery rate (FDR) was determined using Percolator with FDR rate set at 1%.

**SILAC cell culture methods and proteomic sample preparation**

SILAC labeling was performed by growing cells for at least five passages in lysine- and arginine-free SILAC medium (RPMI, Invitrogen) supplemented with 10% dialyzed fetal calf serum (GeminiBio) and 1% Pen/Strep. “Light” and “heavy” media were supplemented with natural lysine and arginine (0.1 mg/mL) for “light”, and 13C-, 15N-labeled lysine and arginine (0.1 mg/mL) for “heavy”, respectively.

**Sample preparation and streptavidin enrichment**

SILAC quantitative proteomics was performed with “heavy” and “light” labeled cells. For K562 cells, SILAC-labeled cells were treated at 1*10^6 cells/mL density in T75 flasks. HeLa and MCF7 cells were grown to confluency and treated in a 10 cm dish. Cells were incubated with DMSO alone (light cells) or chemical probe (25 μM, heavy cells) for 4 hours (for K60P-Bio) or 1 hour (for X4K) in serum-free SILAC RPMI for probe no-probe studies. For inhibitor competition studies, cells were pretreated with inhibitor (KW-2449, 2 μM) for 1 hour before probe treatment of both heavy and light with probe (25 μM) for 2 hours. After incubation, excess probe was removed and supplanted with media for 10 minutes.

Following aspiration of media, cells were collected and then lysed in RIPA lysis buffer (50 mM Tris, 150 mM NaCl, 1% Triton X-100, 0.5% deoxycholate, pH 7.4) supplemented with EDTA-free complete protease inhibitor (Roche) and 1 mM DTT, at 4 °C. After sonication, insoluble debris was cleared by centrifugation (17,000 g, 15 min). BCA assay was performed to normalize Heavy and Light protein concentrations to ~1 mg/ml. Streptavidin agarose beads (50 μL slurry, Pierce) were washed twice with RIPA buffer, and each cell lysate was separately incubated with the beads with rotation overnight at 4 °C. The beads were subsequently washed five times with 0.5 mL of RIPA lysis buffer containing 1 mM DTT, combined together, four times with 0.5 mL PBS, and two times with 2 M Urea in 25 mM ammonium bicarbonate. 500 μL of 6 M Urea in 50 mM ammonium bicarbonate was then added to the beads, and samples were reduced on resin by TCEP (10 mM final) with orbital shaking for 20 minutes at 65℃. Samples were then alkylated by adding iodoacetamide (20 mM final), covered from the light and with orbital shaking, for 40 minutes at 37 °C. The streptavidin agarose beads were collected, washed once with 2 M Urea in 25 mM ammonium bicarbonate, and the buffer exchanged to 2 M Urea in 25 mM ammonium bicarbonate supplemented with 1 mM $CaCl_2$. Enriched proteins were digested on-bead by the incubation of 2 µg sequencing grade trypsin overnight at 37 °C. Following trypsinization, supernatant was collected, acidified with HPLC grade formic acid (2% final, pH 2-3), and peptides were then desalted on ZipTip C18 tips (100 µL, Millipore), dried under vacuum, resuspended with LC-MS grade water (Sigma Aldrich), and then lyophilized. Lyophilized peptides were dissolved in LC-MS/MS Buffer ($H_2O$ with 0.1% formic acid, LC-MS grade, Sigma Aldrich) for proteomic analysis.

**LC-MS/MS Acquisition and Analysis**

Proteomic methods reported are adopted from our previous reports[2]. LC–MS/MS analysis for proteomics samples was performed with an UltiMate 3000 RSLCnano System (Thermo Fisher Scientific) using an Acclaim PepMap RSLC C18 column (75 µm × 15 cm, 2 µm, 100 Å, Thermo Fisher Scientific) with an in-line Acclaim PepMap 100 C18 trap column (75 µm × 2 cm, 3 µm, 100 Å, Thermo Fisher Scientific) heated to 45 °C. The LC system was coupled to an Orbitrap Exploris 480 and Nanospray Flex Ion Source with stainless steel emitter tip (Thermo Fisher Scientific). Mobile phase A was composed of $H_2O$ supplemented with 0.1% formic acid, and mobile phase B was composed of $CH_3CN$ supplemented with 0.1% formic acid. The instrument was run at 0.3 µl min−1 with 2-hour gradients. MS/MS spectra were collected for the entirety of the gradient using a data-dependent, 2-second cycle time setting with the following details: full MS scans were acquired at a resolution of 120,000, scan range of 380 m/z to 1,500 m/z, maximum IT of 25 ms, normalized AGC target of 300% and data collection in profile mode. MS2 scans were performed by high-energy collision dissociation (HCD) fragmentation with a resolution of 15,000, normalized AGC target of 50%, maximum IT of 50 ms, HCD collision energy of 30% and data collection in centroid mode. The isolation window for precursor ions was set to 1.6 m/z. Peptides with a charge state of 1, 7+ and unassigned were excluded, and dynamic exclusion was set to 40 seconds. The RF lens % was set to 40 with a spray voltage value of 2.0 kV and an ionization chamber temperature of 300 °C.

Data were processed using the SEQUEST HT search engine node within the Proteome Discoverer 3.0 software package. Data were searched using a concatenated target/decoy UniProt database of the human proteome with isoforms. Digest enzyme specificity was set to trypsin with up to two missed cleavages allowed, and peptide length was set to between 6 and 144 residues. Precursor mass range was set to 350–6500. Precursor mass tolerance was set to 10 ppm, and fragment mass tolerance was set to 0.02 Da. Up to 4 dynamic modifications were allowed per peptide, including heavy lysine (+8.0142), heavy arginine (+10.0083), oxidized methionine (+15.9949), N-terminal acetylation (+42.0106), N-terminal Met-loss (−131.0405) and N-terminal Met-loss + acetylation (−89.0299). Cysteine carboxyamidomethylation (+57.0215) was set as a static modification. A minimum of two peptides, with a minimum length of 6, was required for protein identification, and false discovery rate (FDR) was determined using Percolator with FDR rate set at 1%. Before quantification, chromatographic alignment was performed, with a maximum retention time difference of 10 min allowed, a mass tolerance of 10 ppm and a minimum signal/noise threshold of 5 required for feature mapping. SILAC ratios were determined using precursor-based quantification in a protein abundance manner based on peak intensity without normalization or scaling using a maximum ratio of 20. Proteins considered as enriched exhibited a probe dependent median SILAC ratio greater than 2 across two biological replicates combined (for Figure 2) and across 4 biological replicates combined (for Figure 3).

#### Supplementary Molecular Modeling Methodology

Visualization of systems was performed using VMD 1.9.4a57[3] or Pymol 2.5 [The PyMOL Molecular Graphics System, Version 2.5 Schrödinger, Inc.]. Analysis was performed with MDTraj 1.9.8[4] and graphs were generated with matplotlib 3.6.2[5].

We initially considered 3 ligands of interest - KW-2449, sunitinib, and XO44 - and the receptors for docking were selected from structures containing these or similar ligands (Figure S1). Crystal structures for sunitinib and XO44 were already available, and crystal structures of axitinib were identified as analogous to KW-2449. The selected structures for docking were 5K9I (Src+XO44), 4WA9 (Abl+axitinib), and 3G0E (KIT+sunitinib). The structures 5K9I and 4WA9 exhibited open binding pockets suitable for docking, the 3G0E binding pocket was much tighter and the A-loop after the DFG phenyl was removed from the 3G0E structure prior to docking.

We determined the analogue space to screen based on examination of the distances between the scaffold and conserved lysine, as well as by examining the sterically similar group on Axitinib. Modifications on the core ring with linker lengths of 0, 1, 2 between the scaffold and phenylsulfonyl fluoride warhead were considered.

Docking experiments were performed with only the attached phenyl ring, and not the fluorosulphonyl group of the warhead. This allowed for the rapid elimination of modification positions that caused steric clashes, whilst minimizing the potential for false negatives due to the inflexibility of the receptor backbone during docking or any clash between the warhead leaving group and lysine. Docking was performed with Smina 2020.12.10[6]. Proteins were prepared for docking using AutoDockTools within the MGLTools software suite 1.5.7[7]. Docking was confined to a box of 30x30x30 Å centered on the binding site. The protein backbone was fixed during docking, but all sidechains facing into the binding pocket were considered flexible. We initially generated 100 poses for each ligand, with the restraint that all poses were > 0.2 Å apart and increased the exhaustiveness of the search to ensure conformations were sampled thoroughly. Poses in which the core-NH was > 2 Å from the crystallographic pose, or that would direct the warhead out of the pocket, were then discarded. Generating multiple poses in multiple receptors was used as the primary indication of probe viability, probes with many generated poses were reasoned to be more likely to form an unstrained covalent bond once the fluorosulfonyl was attached to the ortho-, meta-, or para- positions. Modification positions that did not produce any viable poses, or only produced poses in 1 receptor were eliminated.

The initial template for MD simulations was constructed with CHARMM-GUI[8-10] with the protein from the 4WA9 (chain B, Abl + axitinib) crystal structure. Systems were constructed by replacing the protein in the template system after alignment to its kinase hinge (resid 316 to 318) with proteins from PDB IDs: 4WA9 (chain B), 4TWP (chain B), 2G1T (chain A), 5K9I (chain A). Ligands were aligned to the pharmacophore of axitinib in the template structure. For systems with ligands modified from 5-position of the KW-2449 scaffold (K5*), the initial ligand conformation prior to alignment was adjusted to an alternate rotation of the ring-connecting alkene relative to the axitinib structure (Figure S2). This conformation better directs the warhead towards Lys271 and is necessary to avoid clashes with the curled Abl P-loop. Both alkene rotations engage in the same hydrogen bonds and were observed to freely exchange during simulations, a QM dihedral scan (not shown) predicted both rotations to have within 0.3 kcal/mol energy in solution. For systems of XO44 in proteins from PDB IDs: 4WA9 or 4TWP, the curled P-loop conformations were replaced by alignment with extended conformations from PDB ID: 6BL8[11] to avoid clashes with the ligand. The T315I gatekeeper mutation in 4TWP was reverted to wild-type. Missing residues in the A-loop of 5K9I (chain A) (and residues up to 2 adjacent) were modeled with MODELLER[12]. Minor extensions were modeled for protein termini in Abl so that all proteins were the same length (resid 233-508). Clashing water molecules (< 1 Å from ligand or protein) were shifted to the bulk solvent. All histidines were protonated HSD except His361/375 in Abl and His384/492 in Src which were protonated as HSE.

The linked MD simulations for screening were performed with NAMD 3.0a13[13] for ortho/meta/para warhead positions for all molecules that passed the initial screening and were determined to be synthetically viable: modified at the 5 or 6 position and with a linker length of 0 or 2. Ortho-, meta-, and para- variants were generated for each chosen modification-position and linker combination. The systems were equilibrated, as per CHARMM-GUI NAMD output, with protein heavy atoms restrained using a force constant of 1.0 kcal/mol for the backbone and 0.5 kcal/mol for the sidechain. The ligand core was restrained only by position restraints on the indazole nitrogens, and Lys271 was not restrained. For the linked XO44 simulation the ligand core restraints also included the full ring system except the warhead ring, and the modeled P-loop residues were also unrestrained. The initial NVT equilibration of 250 ps was followed by an additional 1 ns NPT equilibration with the same restraints. Production runs were performed for 100 ns. RMSD calculations for the core displacement, and lysine backbone displacement metrics were calculated on the final 10 ns of simulation frames aligned by all Cα to the 4WA9 crystal structure chain B.

The unlinked simulations to investigate the labelling rate were performed using the CUDA version of Amber 20[14-16]. The systems were converted to amber format using Chamber[17] within ParmEd[18]. The systems were equilibrated, as per CHARMM-GUI Amber output but the ligands were restrained (XO44 heavy atoms, K60P indazole), and modeled protein residues were unrestrained. Production runs were performed for 4 μs.

Force field parameters for each ligand were obtained for the CGenFF 4.4 force field[19, 20] using Paramchem[21, 22]. Parameters were calculated for the unlinked ligand, as well as for a linked form where the fluorine was replaced with a nitrogen and attached lysine sidechain (CB as a methyl group). The bonded terms around the covalent bond were a good match to the model compound (MBSM) used in the development of the CGenFF force field[20], and we reproduced the potential energy scans in MD and QM (not shown). The dihedral parameter between the aromatic and piperazine rings of XO44 selected by Paramchem was a poor analogy and the dihedral 'NG2R62 CG2R64 NG301 CG331' was used instead. The resulting parameters were applied to the unlinked ligand and lysine system as a patch using NAMD's psfgen. The parameters for the charge of the warhead fluorine and S-F bond length were incorrect and were re-fit. The charges of the warhead and adjacent phenyl ring were fit using FFParam[23] according to the CGenFF parameterization guidelines, using Gaussian[24] and OpenMM[25] for the required QM and MD calculations respectively. The S-F bond-length was set to 1.62 Å as per the QM optimization.

#### Supplementary Molecular Modeling Discussion

When conducting MD simulations of the unlinked ligand to investigate the labelling rate, the simulations were performed using protein conformations taken from X-ray structures of the kinase co-crystallized with axitinib (Abl) and XO44 (Src): Abl DFG-out (Abl/DFG-out, 4WA9), Abl DFG-in (Abl/DFG-in, 4TWP), Src DFG-in (Src/DFG-in, 5K9I), and Abl DFG-in/αC-helix-out (Abl/DFG-in/αC-helix-out, 2G1T). This resulted in a total of 2 duplicates of 4 systems for each of the 2 ligands (16 × 4 µs simulations).

The DFG-out simulations for both ligands had the lowest sampling of reaction-ready proximities. In this conformation, π-stacking interactions between the warhead ring and the phenylalanine sidechain of the DFG motif effectively displaced the warhead, disrupting the core hydrogen bonds and fencing it away from the target Lysine (Figure 2C, 2D). This suggests that both ligands have a preference for reversible binding to the DFG-in conformation of Abl kinase.

In the Abl DFG-in simulations XO44 outperformed K60P in spite of unstable interactions with the Abl P-loop. The XO44 warhead was observed to flip into and out of the binding pocket, resulting in closer proximities than in Src DFG-in where the warhead position remained stable. In 1 replicate of each DFG-in Abl-XO44, the P-loop curled up preventing the warheads re-entry. K60P was stable in the DFG-in simulations, but we observed that the warhead ring tended to cause the Lysine to be displaced away from the warhead towards the P-loop. The Lys to DFG-in-Asp salt-bridge strained to form over the top of the ligand, inducing the scaffold to angle slightly deeper into the cleft than our predicted poses and more frequently adopt the 180° rotated alkene position (Figure S2).

All K60P simulations resulted in the αC-helix adopting the out conformation after a few 100 ns, likely due to charge-repulsion from the warhead, thus preventing the formation of a salt-bridge between the Lysine and αC-helix-Glu. For the XO44 simulations the αC-helix remained in its initial conformation. The XO44 crystal structures are in the αC-helix-out conformation, but it is unclear if this only occurs post-reaction.

#### Supplementary Kinetic Modeling Methodology

To characterize the kinetics of ABL1-probe binding, the experimental data were fit with numerical simulations of irreversible kinetic schemes. The basic kinetic scheme underlying all systems is defined with two-step inactivation and one-step nonspecific binding (Scheme 1). The two-step inactivation comprises a reversible non-covalent association/dissociation step followed by an irreversible covalent reaction step. The protein species that exists solely for non-specific binding, $P_{ns}$, is unconsumed to approximate the slow saturation of nonspecific binding sites.

Ordinary differential equations (ODEs) were constructed for each transition in the scheme and integrated together numerically to yield the time evolution of each species concentration. These simulations used the same basic conditions as the experiments. The two independent free protein species, P and $P_{ns}$, had an initial concentration of 0.357 μM. Time-responses used an initial probe concentration of 10 μM. Dose-responses used a timepoint of one hour.

The time- and dose-responses on their own do not offer much value for fitting unnormalized labeling data, but they complement each other well when used simultaneously. The fitting procedure, which is done for each probe individually, leverages this idea by considering the residuals between all corresponding experimental and simulated data at once. The simulations are parameterized with the rate constants for each kinetic transition and a normalizing factor (λ) for each experimental data set. Therefore, a reasonable set of rate constants must yield a reasonable fit to every replicate of both time- and dose-responses. The simulated response is 1) the sum of the concentrations of the specifically- and nonspecifically-bound covalent complexes, [P-L] and [$P_{ns}$-L], 2) normalized by the initial protein concentration, and 3) multiplied by the λ parameter for the corresponding experiment. Note that when plotting and analyzing the outcome of the fitting procedure, the simulated response is left normalized to initial protein concentration, and the experimental datapoints are divided by λ as an effective normalization. The residuals were minimized using LMFIT[26] with the Levenberg-Marquardt least-squares (LS) algorithm[27] and Trust Region Reflective method[28].

Initial guesses and bounds of $k_{on}$ and $k_{off}$ were informed by a comprehensive kinetic survey between kinases and competitive ligands[29]. To efficiently focus the sampling on the binding affinity, ($K\text{d}=k_{off}/k_{on}$), with minimal degeneracy, either $k_{on}$ or $k_{off}$ must be fixed. Fixing $k_{on}$ is more ideal, since it is largely diffusion limited and differences in the ligands are more likely to be represented by differences in the $k_{off}$. This was a viable strategy for K50P and K60P, but X4K required a fixed $k_{off}$. X4K was possibly rate-limited by $k_{on}$, so the rest of the kinetic parameters and affinity would be largely dependent on the fixed value of $k_{on}$. The $k_{on}$ was fixed for the KW probes at 100 times less than the value for KW-449/ABL1, $1.82\times10^{-2}$ $\mu M^{-1}s^{-1}$. For X4K, $k_{off}$ was fixed according to analogue VX-680, $8.95\times10^{-2}$ $s^{-1}$. We discarded the 2- and 60-minute timepoints in the X4K time-response experiment, because no reasonable fits to the dose-response could be produced otherwise.

The best parameters from the LS minimization were used to initialize Metropolis Monte-Carlo (BMMC) sampling, which used the same fitting procedure to calculate the chi-square value from the residuals as an "energy" for the purposes of acceptance. The normalization factors were held constant in addition to the parameters that were already fixed in the LS fit. The step size for each variable parameter was chosen uniformly between +/- 5% of the respective best-fit value. To reduce the effect of autocorrelation, only every 500th step in the BMMC chain was kept. The presented range of each variable parameter is the highest density interval for 95% of its BMMC distribution[30].

Although the absolute scale of labeling data with respect to total protein concentration is unknown, the relative scaling between the KW probes was approximated as an inter-ligand scaling factor from the one time-response (Bio Rep 1) and two dose-responses (Bio Reps 3 and 4) that captured K50P and K60P together on the same gel. This inter-ligand scaling factor for each replicate was calculated as the K60P:K50P ratio of the final response. Probe K60P was fit independently, yielding a standard error for each normalization factor that was calculated as the square root of the product of the diagonal elements of the covariance matrix and the reduced chi-square. The initial guess of the respective K50P normalization factor was the K60P best fit multiplied by the inter-ligand scalar. The bounds were the initial guess +/- two scaled standard errors. The propagation of uncertainty and the flexibility of inter-ligand scaling, though crude, enabled the K50P fitting to utilize the relative scaling observed in all three Bio Reps.

Following the observations of relative scaling between the KW probes, the kinetic scheme of K50P was modified to account for sub-stoichiometry. Two modified versions of the scheme were created to allow the fraction of specific binding to plateau below 100% of the initial protein concentration, $Mod_{del}$ (Scheme S1) and $Mod_{sil}$ (Scheme S2). $Mod_{del}$ introduces the parameter $k_{del}$, *i.e.* the rate constant of K50P "deletion" (del) and models a ligand-limited plateau. $Mod_{sil}$ introduces the parameter $Frac_{sil}$, *i.e.* the fraction of "silent" (sil) ligand and models a protein-limited plateau. The term "silent" stems from the scenario where a portion of the total ligand population is consuming protein without contributing to the observable. Note that the measured effect of $Mod_{sil}$ would be equivalent had the silent population been protein instead, effectively modeling the scenario where the protein is the affected species, *e.g.* if the ligand induces and traps a silencing and rapidly-equilibrating conformational change in the protein. The sub-stoichiometric modeling is elaborated upon in the supplementary kinetic modeling discussion.

Scheme S1. Additional transition to express sub-stoichiometric covalent occupancy in terms of ligand "deletion".

$$\mathrm{L} \xrightarrow{k_{\mathrm{del}}} 0\mathrm{L}$$

Scheme S2. Additional transitions to express sub-stoichiometric covalent occupancy in terms of a "silent" ligand population.

$$\mathrm{P} + \mathrm{L_{sil}} \underset{k_{\mathrm{off}}}{\overset{k_{\mathrm{on}}}{\rightleftharpoons}} \mathrm{PL_{sil}} \xrightarrow{k_{\mathrm{inact}}} \mathrm{P\text{-}L_{sil}}$$

$$\mathrm{P_{ns}} + \mathrm{L_{sil}} \xrightarrow{k_{\mathrm{ns}}} \mathrm{P_{ns}} + \mathrm{P_{ns}\text{-}L_{sil}}$$

Note that the original ligand concentration is distributed over two populations, L and $L_{sil}$, where the proportion of silent ligand is defined as the parameter, $Frac_{sil}$.

#### Supplementary Kinetic Modeling Discussion

Early in the study, the experimental time-dependent data displayed a striking difference in unnormalized intensity between K50P and K60P. By the six-hour endpoint, K60P exceeded twice the intensity of K50P, which had reached an apparent plateau. Since the specific binding of both irreversible probes should plateau at 100% of the protein concentration, we reasoned that the difference in intensity could be caused by greater nonspecific binding by K60P or an unknown mechanism causing K50P to plateau sub-stoichiometrically. Reasonable estimates of the probes' nonspecific-binding rate constants could be fitted to the dose-responses. Yet, the nonspecific binding only accounted for about half of the difference in probe intensity. The remaining half was achieved by inducing either ligand-limited or protein-limited sub-stoichiometry with one of two generic modifications, $Mod_{del}$ or $Mod_{sil}$, to the base kinetic model of K50P. $Mod_{del}$ introduces a one-step transition that deletes (del) the ligand and lowers its effective concentration over time (Scheme S1). $Mod_{sil}$ introduces a silent (sil) fraction of the ligand population that can covalently bind protein without producing signal (Scheme S2), akin to the transducer ratio for the operational model of agonism[31]. Both modifications caused the same apparent behavior (Figure S9).

The models of sub-stoichiometry and their associated parameters are not representative of any specific mechanism. They instead represent generic mechanisms that act on the simulated response to align with empirical observations and cover a broad range of possibilities. For example, $Mod_{sil}$ covers covalent unbinding within the quenching period of the assay, and $Mod_{del}$ covers hydrolysis of the sulfonylfluoride warhead. An important purpose of these modifications is to demonstrate how estimations of canonical kinetic features, such as the affinity and the rate constant of inactivation, are dependent on the model of sub-stoichiometry. The fitted values of $K_d$ and $k_{inact}$ were mostly unaffected when the kinetic analysis included $Mod_{sil}$ but were approximately halved when it included $Mod_{del}$. These features are especially susceptible to complex versions of $Mod_{del}$. As an illustrative yet unlikely example, modeling depletion of productive ligand via protein-mediated hydrolysis at the binding site lowers the estimated $K_d$ by about an order of magnitude, since the complete depletion within the experimental timeframe depends on the quality of reversible binding.

In any case, the inactivation rate constants, $k_{inact}$, for the KW ligands were consistent with previously reported values for XO44 analogues ($k_{obs}$ ranging from $3 \times 10^{-4}$ to $8 \times 10^{-4}$ $s^{-1}$)[32] and to a similarly positioned probe (ABP4) binding to ABL1 ($k_{inact} = 0.778 \times 10^{-4}$ $s^{-1}$)[33]. The median $k_{inact}$ values for $K50P_{sil}$ and K60P were nearly identical despite their empirical differences. However, the median nonspecific-binding rate constant, $k_{ns}$, for K60P was about three times larger than that of both K50P variants, consistent with the qualitative trends observed in nonspecific reactivity assays (Figure S11).

Unlike the KW probes, the $k_{inact}$ of X4K could be fast enough to compete with $k_{on}$, (and the fixed $k_{off}$) and prevent an effective equilibrium of association and dissociation. Consequently, the apparent $K_d$ is less meaningful, particularly in the commonly-sampled space where $k_{inact}$ greatly exceeds $k_{off}$. It is also possible that the $k_{on}$ is entirely rate-limiting, in which case the upper bound of $k_{inact}$ is indeterminable from the labeling data. Yet, this is not the only factor of ambiguity surrounding the X4K analysis. In the dose-response, X4K appeared to covalently bind 100% of ABL1 within the incubation time at doses too low to rapidly saturate the reversibly associated state and yield $k_{on} \times [L] >> k_{inact}$. Therefore, unlike the KW probes, the unique role of $k_{inact}$ in modulating the saturation-driven plateau of covalent occupancy could not be leveraged for fitting $k_{inact}$[34]. With only the $EC_{50}$ to modulate, the $k_{inact}$ and $k_{on}$ were highly correlative in the fitting. In turn, the sampled values of $k_{inact}$ were often also dependent on the assumed value of $k_{off}$

($8.95'10^{-2}$ $s^{-1}$). Still, as pointed out in the main text, it is clear qualitatively that the X4K labeling is faster than that of K50P and K60P due to a faster $k_{inact}$. This increased reactivity is also reflected in the estimated nonspecific rate constant $k_{ns}$, which has a median approximately three times greater for X4K than K60P.

#### Supplementary Synthetic methods and characterization

##### General Procedures

All reactions were carried out under a nitrogen atmosphere with dry solvents under anhydrous conditions, unless otherwise noted. Super-dry solvents (water ≤ 30–50 ppm) including tetrahydrofuran (THF), toluene (PhMe), *N,N*-dimethylformamide (DMF), ethyl ether ($Et_2O$) and dioxane were purchased from Thermo Fisher and used directly. Deionized water was used throughout this work. Reagents were purchased at the highest commercial quality and used without further purification, unless otherwise stated. Solvents for chromatography were used as supplied by Thermo Fisher. Reactions were monitored by thin layer chromatography (TLC) carried out on 0.24 ± 0.03 mm Merck silica gel plates (GF254) using UV light as visualizing agent and aqueous ammonium cerium nitrate/ammonium molybdate, aqueous phosphomolybdic acid or basic aqueous potassium permanganate as developing agent. Merck silica gel (particle size: 40–63 µm, pore size: 60 Å) was used for flash column chromatography. NMR spectra were recorded on Bruker Avance III HD nanobay 400 MHz and Bruker Avance Neo 500 MHz instruments. The following abbreviations are used to designate multiplicities: s = singlet, d = doublet, t = triplet, q = quartet, m = multiplet, quint = quintet, br = broad. Accurate mass measurements and final probe purification were obtained and performed using an Agilent 1290 Infinity II with an Agilent 5 Prep C18 50 × 21.2mm column.

##### Experimental procedures

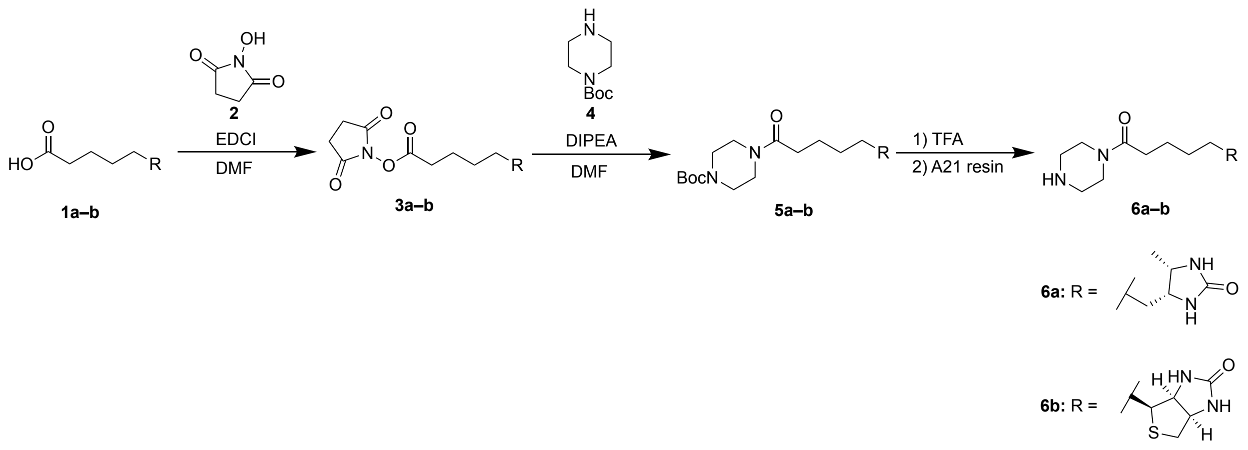

**Compound 3a**: To a stirred solution of D-desthiobiotin **1a** (0.53 g, 2.5 mmol) and *N*-hydroxysuccinimide **2** (0.43 g, 3.75 mmol) in DMF (8.4 mL) was added EDCI (0.72 g, 3.75 mmol). The reaction mixture was allowed to stir at room temperature overnight before removing the solvent under vacuum. Water (8.4 mL) was added to the residue. The precipitate was collected by filtration to give compound **3a** (0.78 g, 95%) as a white solid, which was used in the next step without further purification. $^1$H NMR (500 MHz, DMSO-$d_6$) δ 6.28 (s, 1H), 6.10 (s, 1H), 3.59 (p, $J$ = 6.7 Hz, 1H), 3.47 (q, $J$ = 7.4 Hz, 1H), 2.79 (s, 4H), 2.65 (t, $J$ = 7.3 Hz, 2H), 1.60 (p, $J$ = 7.1 Hz, 2H), 1.46 – 1.09 (m, 6H), 0.94 (d, $J$ = 6.4 Hz, 3H).

**Compound 3b** was synthesized using the same method as **3a**. D-biotin **1b** (0.50 g, 2.04 mmol) was used instead of D-desthiobiotin **1a** to give compound **3b** (0.63 g, 90%) as a white solid, which was used in the next step without further purification. $^1$H NMR (400 MHz, DMSO-$d_6$) δ 6.42 (s, 1H), 6.36 (s, 1H), 4.31 (t, $J$ = 6.4 Hz, 1H), 4.19 – 4.12 (m, 1H), 3.11 (q, $J$ = 6.6 Hz, 1H), 2.92 – 2.81 (m, 1H), 2.82 (s, 4H), 2.76 – 2.63 (m, 2H), 2.59 (d, $J$ = 12.4 Hz, 1H), 1.66 (p, $J$ = 7.1 Hz, 3H), 1.57 – 1.35 (m, 3H).

**Compound 5a**: To stirred solution of **3a** (0.48 g, 1.5 mmol) and 1-boc-piperazine **4** (0.55 g, 3.0 mmol) in DMF (7.7 mL) was added DIPEA (0.52 mL, 4.5 mmol). The reaction mixture was allowed to stir at room temperature overnight before removing the solvent under vacuum. Water (10 mL) was added to the residue, the participate was filtered and washed with water (5.0 mL) and $Et_2O$ (5.0 mL) to give compound **5a** (0.45 g, 76%) as a white solid. The compound was used in the next step without further purification. $^1$H NMR (500 MHz, DMSO-$d_6$) δ 6.30 (s, 1H), 6.11 (s, 1H), 3.58 (p, $J$ = 6.7 Hz, 1H), 3.46 (t, $J$ = 6.7 Hz, 1H), 3.42 – 3.36 (m, 4H), 3.25 (s, 2H), 2.28 (t, $J$ = 7.5 Hz, 2H), 1.46 (q, $J$ = 7.4 Hz, 2H), 1.39 (s, 9H),

1.35 – 1.12 (m, 6H), 0.94 (d, *J* = 6.3 Hz, 3H).

**Compound 5b** was synthesized using the same method as **5a**. **3b** (0.48 g, 1.4 mmol) was used instead of **3a** to give compound **5b** (0.45 g, 76%) as a white solid, which was used in the next step without further purification. $^{1}$H NMR (400 MHz, $CDCl_3$) δ 5.34 (s, 1H), 4.76 (s, 1H), 4.55 – 4.48 (m, 1H), 4.33 (ddd, J = 7.8, 4.6, 1.6 Hz, 1H), 3.59 (d, J = 4.3 Hz, 2H), 3.42 (m, 6H), 3.18 (td, J = 7.3, 4.5 Hz, 1H), 2.93 (dd, J = 12.8, 5.0 Hz, 1H), 2.73 (d, J = 12.8 Hz, 1H), 2.37 (t, J = 7.3 Hz, 2H), 1.82 – 1.63 (m, 4H), 1.47 (s, 11H).

**Compound 6a**: A solution of **5a** (0.40 g, 1.0 mmol) in ice-cold TFA (5.0 mL) was slowly warmed up to room temperature and stirred at that temperature for 1h before removing the solvent under vacuum. The residue was co-evaporated with EtOAc (2 × 10 mL) and MeOH (2 × 10 mL) before it was dissolved in MeOH (5.0 mL). The solvent was evaporated under vacuum and ether was added to precipitate compound **6a** (0.37 g, 93%) as a white solid. $^{1}$H NMR (400 MHz, DMSO-$d_6$) δ 8.61 (s, 2H), 6.31 (s, 1H), 6.13 (s, 1H), 3.70 – 3.56 (m, 5H), 3.53 – 3.44 (m, 1H), 3.14 – 2.97 (m, 4H), 2.33 (t, *J* = 7.5 Hz, 2H), 1.49 (p, *J* = 7.2 Hz, 2H), 1.41 – 1.14 (m, 6H), 0.97 (d, *J* = 6.4 Hz, 3H).

**Compound 6b** was synthesized using the same method as **6a**. **5b** (0.40 g, 0.97 mmol) was used instead of **5a** to give compound **6b** (0.37 g, 93%) as a white solid, which was used in the next step without further purification. $^{1}$H NMR (400 MHz, DMSO-$d_6$) δ 8.66 (s, 2H), 6.42 (t, *J* = 1.8 Hz, 1H), 6.37 (s, 1H), 4.32 (ddt, *J* = 7.6, 5.3, 1.2 Hz, 1H), 4.14 (ddd, *J* = 7.7, 4.4, 1.8 Hz, 1H), 3.62 (p, *J* = 3.8 Hz, 4H), 3.15 – 2.99 (m, 5H), 2.83 (dd, *J* = 12.4, 5.1 Hz, 1H), 2.63 – 2.53 (m, 1H), 2.39 – 2.28 (m, 2H), 1.70 – 1.26 (m, 6H).

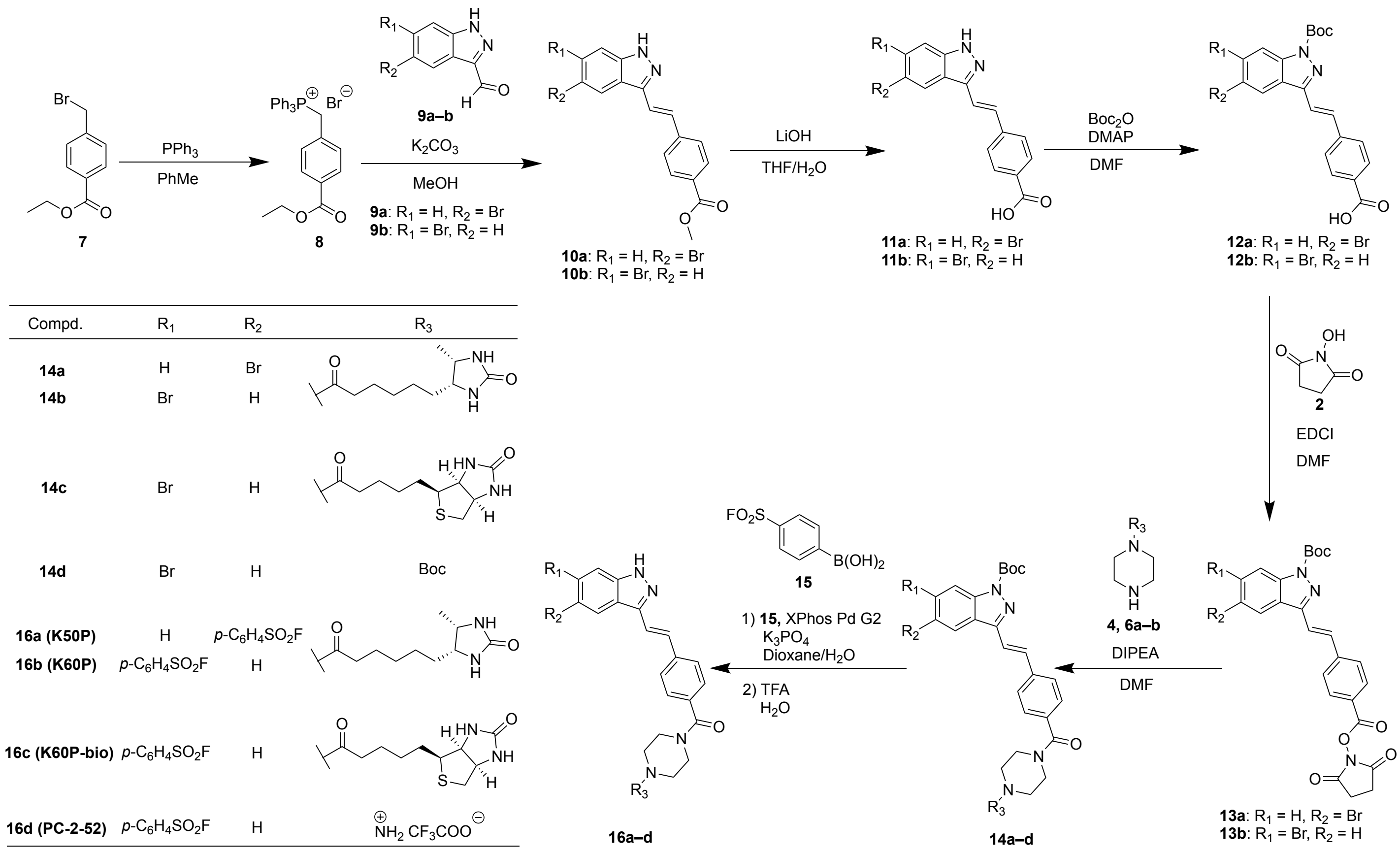

**Compound 8**: To a stirred solution of ethyl 4-(bromomethyl)benzoate **7** (6.4 g, 26 mmol) in PhMe (25 mL) was added $PPh_3$ (7.5 g, 29 mmol). The mixture was heated to 100 °C and allowed to stir at that temperature overnight before it was filtered and washed with ethyl ether (30 mL). The solid was dried over vacuum to give compound **8** (13 g, 98%) as a white solid. $^1$H NMR (400 MHz, DMSO-$d_6$) δ 8.04 – 7.86 (m, 3H), 7.86 – 7.62 (m, 14H), 7.13 (dd, J = 8.5, 2.5 Hz, 2H), 5.30 (d, J = 16.3 Hz, 2H), 4.28 (q, J = 7.1 Hz, 2H), 1.30 (t, J = 7.1 Hz, 3H).

**Compound 10a**: To a stirred solution of **8** (6.9 g, 14 mmol) and 5-bromo-1H-indazole-3-carbaldehyde **9a** (2.6 g, 12 mmol) in MeOH (19 mL) was added $K_2CO_3$ (4.7 g, 34 mmol). The reaction mixture was allowed to stir at room temperature overnight before it was cooled down to 0 °C and acidified with conc. HCl dropwise until fully precipitated. The resultant mixture was filtered and washed with MeOH (10 mL) and water (10 mL). The solid was dried under vacuum to give **10a** (0.67 g, 18%) as a beige solid. $^1$H NMR (400 MHz, DMSO-$d_6$) δ 8.47 (s, 1H), 7.98 (d, *J* = 8.4 Hz, 2H), 7.89 (d, *J* = 8.4 Hz, 2H), 7.75 (d, *J* = 16.7 Hz, 1H), 7.61 – 7.52 (m, 2H), 7.49 (dd, *J* = 8.7, 1.8 Hz, 1H), 3.87 (s, 3H).

**Compound 10b** was synthesized using the same method as **10a**. 6-bromo-1H-indazole-3-carbaldehyde **9b** (1.3 g, 5.8 mmol) was used instead of **9a**, with the same equivalents of **8** and $K_2CO_3$ to give compound **10b** (0.19 g, 10%) as a beige solid. $^1$H NMR (400 MHz, DMSO-$d_6$) δ 8.20 (d, $J$ = 8.7 Hz, 1H), 7.98 (d, $J$ = 8.0 Hz, 2H), 7.87 (d, $J$ = 8.1 Hz, 2H), 7.80 (s, 1H), 7.72 (d, $J$ = 16.7 Hz, 1H), 7.58 (d, $J$ = 16.7 Hz, H), 7.38 – 7.31 (m, 1H), 3.87 (s, 3H).

**Compound 11a**: To a stirred solution of **10a** (0.67 g, 1.9 mmol) in THF (19 mL) was added 4.0 M LiOH (0.46 g, 19 mmol) aqueous solution (4.7 mL). The reaction mixture was warmed up to 40 °C and allowed to stir at that temperature overnight. 12N HCl was added dropwise till pH was ~4 which resulted in the precipitation of a beige solid. The precipitate was filtered and rinsed with water to get **11a** (0.52g, 81% yield). $^1$H NMR (400 MHz, DMSO-$d_6$) δ 13.45 (s, 1H), 12.91 (s, 1H), 8.50 (d, $J$ = 1.7 Hz, 1H), 7.96 (d, $J$ = 8.2 Hz, 2H), 7.87 (d, $J$ = 8.2 Hz, 2H), 7.73 (d, $J$ = 16.7 Hz, 1H), 7.59 (d, $J$ = 16.7 Hz, 1H), 7.56 – 7.52 (m, 2H).

**Compound 11b** was synthesized using the same method. **10b** (0.19 g, 0.53 mmol) was used instead of **10a**, with the same equivalents of LiOH to give compound **11b** (0.15 g, 82%) as a beige solid. $^1$H NMR (400 MHz, DMSO-$d_6$) δ 13.62 (s, 1H), 8.20 (d, $J$ = 8.7 Hz, 1H), 7.95 (d, $J$ = 8.1 Hz, 2H), 7.88 – 7.79 (m, 3H), 7.70 (d, $J$ = 16.7 Hz, 1H), 7.58 (d, $J$ = 16.7 Hz, 1H), 7.34 (dd, $J$ = 8.6, 1.6 Hz, 1H).

**Compound 12a**: To a stirred solution of **11a** (0.26 g, 0.76 mmol) and $Boc_2O$ (0.35 g, 0.37 mL, 1.6 mmol) in DMF (3.0 mL) was added DMAP (36 mg, 0.30 mmol). The reaction mixture was allowed to stir at room temperature overnight before the solvent was removed under vacuum. The residue was dissolved in EtOAc (15 mL) and washed with saturated aq. $NaHCO_3$ (15 mL), brine (2 × 10 mL) dried over anhydrous $Na_2SO_4$, and filtered. The solvent was evaporated under vacuum to give **12a** (0.23 g, 68%) as a light yellow solid. $^1$H NMR (400 MHz, DMSO-$d_6$) δ 12.99 (s, 1H), 8.65 (d, $J$ = 1.8 Hz, 1H), 8.08 (d, $J$ = 8.9 Hz, 1H), 8.02 – 7.94 (m, 4H), 7.86 – 7.76 (m, 3H), 1.68 (s, 9H).

**Compound 12b** was synthesized using the same method as **12a**. **11b** (0.15 g, 0.44 mmol) was used instead of **11a**, with the same equivalents of $Boc_2O$ and DMAP to give compound **12b** (0.18 g, 90%) as a light yellow solid. $^1$H NMR (500 MHz, DMSO-$d_6$) δ 8.35 – 8.27 (m, 2H), 7.96 – 7.80 (m, 4H), 7.76 – 7.68 (m, 1H), 7.68 – 7.60 (m, 2H), 1.65 (s, 9H).

**Compound 13a**: To a stirred solution of **12a** (0.23 g, 0.52 mmol) and *N*-hydroxysuccinimide **2** (89 mg, 0.78 mmol) in DMF (2.6 mL) was added EDCI (0.15 g, 0.78 mmol). The reaction mixture was allowed to stir at room temperature overnight before it was diluted with DCM (5.0 mL). The resultant mixture was washed with water (2 × 10 mL), brine (2 × 10 mL), dried over anhydrous $Na_2SO_4$, and filtered. The solvent was evaporated under vacuum to give **13a** (0.19 g, 69%) as a light yellow solid. $^1$H NMR (400 MHz, DMSO-$d_6$) δ 8.68 (d, *J* = 1.9 Hz, 1H), 8.17 – 8.11 (m, 4H), 8.11 – 8.06 (m, 1H), 7.87 (d, *J* = 4.3 Hz, 2H), 7.85 – 7.80 (m, 1H), 2.92 (s, 4H), 1.68 (s, 9H).

**Compound 13b** was synthesized using the same method. **12b** (0.18 g, 0.41 mmol) was used instead of **12a**, with the same equivalents of *N*-hydroxysuccinimide **2** and EDCI to give **13b** (0.15 g, 70%) as a light yellow solid. $^1$H NMR (400 MHz, DMSO-$d_6$) δ 8.39 (d, *J* = 8.6 Hz, 1H), 8.33 (d, *J* = 1.7 Hz, 1H), 8.12 (q, *J* = 8.7 Hz, 4H), 7.86 (s, 2H), 7.68 (dd, *J* = 8.5, 1.7 Hz, 1H), 2.92 (s, 4H), 1.68 (s, 9H).

**Compound 14a**: To a stirred solution of **13a** (0.19 g, 0.35 mmol) and **6a** (0.17 g, 0.42 mmol) in DMF (1.8 mL) was added DIPEA (0.14 g, 0.19 mL, 1.1 mmol). The reaction mixture was allowed to stir at room temperature for 4h before the solvent was removed under vacuum. The residue was precipitated with water (2.0 mL), and the solid was collected by filtration followed by drying over vacuum to give **14a** (0.20 g, 79%) as a beige powder. $^1$H NMR (400 MHz, DMSO-$d_6$) δ 8.71 – 8.60 (m, 1H), 8.11 – 7.89 (m, 4H), 7.87 – 7.71 (m, 3H), 7.49 (d, *J* = 8.0 Hz, 1H), 6.31 (s, 1H), 6.12 (s, 1H), 3.65 – 3.40 (m, 8H), 2.33 (s, 2H), 1.68 (s, 9H), 1.50 (brs, 2H), 1.43 – 1.18 (m, 8H), 0.96 (d, J = 6.3 Hz, 3H).

**Compound 14b** was synthesized using the same method as **14a**. **13b** (75 mg, 0.14 mmol) was used instead of **13a**, with the same equivalents of DIPEA to give **14b** (85 mg, 86%) as a beige powder as a mixture of isomers which was used without further purification. $^1$H NMR (400 MHz, DMSO-$d_6$) δ 8.38 – 8.22 (m, 1H), 8.08 – 7.86 (m, 2H), 7.84 – 7.63 (m, 3H), 7.57 – 7.24 (m, 3H), 6.31 (s, 1H), 6.12 (s, 1H), 3.60 (q, *J* = 6.7 Hz, 8H), 2.32 (s, 2H), 1.56 – 1.13 (m, 10H), 0.96 (d, *J* = 6.3 Hz, 3H).

**Compound 14c** was synthesized using the same method as **14a**. **13b** (0.18 g, 0.33 mmol) and **6b** (0.13 g, 0.40 mmol) were used instead of **13a** and **6a**, with the same equivalents of DIPEA to give **14c** (0.16 g, 63%) as a beige powder. $^1$H NMR (400 MHz, DMSO-$d_6$) δ 8.71 – 8.60 (m, 1H), 8.12 – 8.02 (m, 2H), 8.00 – 7.90 (m, 2H), 7.86 – 7.70 (m, 3H), 7.49 (d, *J* = 8.1 Hz, 1H), 6.43 (s, 1H), 6.36 (s, 1H), 4.31 (t, *J* = 6.5 Hz, 1H), 4.14 (s, 1H), 3.67 – 3.35 (m, 6H), 3.11 (s, 1H), 2.83 (dd, *J* = 12.5, 5.3 Hz, 1H), 2.68 (s,1H), 2.59 (d, *J* = 11.9 Hz, 1H), 2.34 (s, 2H), 1.68 (s, 9H), 1.57 – 1.18 (m, 7H).

**Compound 14d** was synthesized using the same method as **14a**. **13b** (0.18 g, 0.33 mmol) and **4** (72 mg, 0.40 mmol) were used instead of **13a** and **6a**, with the same equivalents of DIPEA to give **14d** (0.16 g, 80%) as a beige powder. $^1$H NMR (400 MHz, DMSO-$d_6$) δ 8.41 – 8.28 (m, 2H), 8.04 (d, *J* = 2.3 Hz, 1H), 7.93 – 7.62 (m, 5H), 7.48 (d, *J* = 8.1 Hz, 1H), 3.63 – 3.34 (m, 8H), 1.68 (s, 9H), 1.41 (dd, *J* = 4.0, 2.0 Hz,

9H).

**Compound 16a (K50P)**: To a stirred solution of **14a** (50 mg, 71 $\mu$mol), **15** (22 mg, 0.11 mmol)[35] and $K_3PO_4$ (45 mg, 0.21 mmol) in dioxane (3.5 mL) and water (0.35 mL) was added XPhos Pd G2 (5.5 mg, 10 mol%). The reaction mixture was heated up to 40 °C and allowed to stir at that temperature overnight before it was diluted with EtOAc (5.0 mL) after cooling to room temperature. The resultant mixture was allowed to pass through a pad of Celite®, washed with water (10 mL), dried over anhydrous $Na_2SO_4$, and filtered. The solvent was evaporated under vacuum and the residue was dissolved in TFA (2.0 mL) and water (2.0 mL). The reaction mixture was allowed to stir at room temperature for 30 min before the solvent was removed under vacuum. The residue was purified by prep-HPLC with ACN:water (30:70 to 80:20) to give **16a** (**K50P**) (9.0 mg, 19%) as a pale yellow solid. $^1$H NMR (400 MHz, DMSO-$d_6$) δ 13.43 (s, 1H), 8.65 (d, $J$ = 1.7 Hz, 1H), 8.23 (s, 4H), 7.89 – 7.82 (m, 3H), 7.79 – 7.63 (m, 3H), 7.48 (d, $J$ = 8.1 Hz, 2H), 6.31 (s, 1H), 6.12 (s, 1H), 3.71 – 3.47 (m, 8H), 2.34 (d, $J$ = 9.6 Hz, 2H), 1.59 – 1.05 (m, 10H), 0.97 (d, $J$ = 6.4 Hz, 3H). $^{19}$F NMR (500 MHz, DMSO-$d_6$) δ 67.00. LC-MS (ESI-API): calculated for $C_{36}H_{40}FN_6O_5S^+$ [M+H]$^+$: 687.3, found: 687.3.

**Compound 16b (K60P)** was synthesized using the same method as **16a**. **14b** (85 mg, 0.12 mmol) was used of instead of **14a**, with the same equivalents of **15**, $K_3PO_4$ and XPhos Pd G2 to give **16b** (8.0 mg, 10%)] as a pale yellow solid. $^1$H NMR (400 MHz, DMSO-$d_6$) δ 13.49 (s, 1H), 8.39 (d, J = 8.5 Hz, 1H), 8.28 – 8.17 (m, 4H), 7.95 (d, J = 1.5 Hz, 1H), 7.84 (d, J = 8.1 Hz, 2H), 7.74 – 7.58 (m, 3H), 7.48 (d, J = 8.1 Hz, 2H), 6.32 (s, 1H), 6.12 (s, 1H), 3.71 – 3.46 (m, 8H), 2.34 (d, J = 11.1 Hz, 2H), 1.58 – 1.07 (m, 10H), 0.97 (d, J = 6.4 Hz, 3H). $^{19}$F NMR (500 MHz, DMSO-$d_6$) δ 66.81. LC-MS (ESI-API): calculated for $C_{36}H_{40}FN_6O_5S^+$ [M+H]$^+$: 687.3, found: 687.2.

**Compound 16c (K60P-Bio)** was synthesized using the same method as **16a**. **14c** (0.12 g, 0.16 mmol) was used instead of **14a**, with the same equivalents of **15**, $K_3PO_4$ and XPhos Pd G2 to give **16c** (**K60P-bio**) (8.5 mg, 7%)] as a pale yellow solid. $^1$H NMR (400 MHz, DMSO-$d_6$) δ 13.41 (s, 1H), 8.31 (d, $J$ = 8.6 Hz, 1H), 8.21 – 8.09 (m, 4H), 7.87 (s, 1H), 7.76 (d, $J$ = 8.0 Hz, 2H), 7.67 – 7.50 (m, 3H), 7.40 (d, $J$ = 8.0 Hz, 2H), 6.35 (s, 1H), 6.28 (s, 1H), 4.32 – 4.19 (m, 1H), 4.07 (s, 1H), 3.66 – 3.35 (m, 8H), 3.12 – 2.96 (m, 1H), 2.76 (dd, $J$ = 12.4, 5.0 Hz, 1H), 2.51 (d, $J$ = 12.3 Hz, 1H), 2.26 (m, 2H), 1.63 – 1.21 (m, 6H). $^{19}$F NMR (500 MHz, DMSO-$d_6$) δ 66.86. LC-MS (ESI-API): calculated for $C_{36}H_{38}FN_6O_5S_2^+$ [M+H]$^+$: 717.2, found: 717.2.

**Compound 16d (PC-2-52)** was synthesized using the same method as **16a**. **14d** (40 mg, 66 $\mu$mol) was used instead of **14a**, with the same equivalents of **15**, $K_3PO_4$ and XPhos Pd G2 to give **16d** (**PC-2-52**) (4.9 mg, 15%) as its TFA salt as a pale yellow solid. $^1$H NMR (400 MHz, DMSO-$d_6$) δ 13.51 (s, 1H), 8.88 (s, 2H), 8.39 (d, $J$ = 8.5 Hz, 1H), 8.30 – 8.16 (m, 4H), 7.95 (d, $J$ = 1.5 Hz, 1H), 7.91 – 7.81 (m, 2H), 7.77 – 7.58 (m, 3H), 7.58 – 7.44 (m, 2H), 3.72 (s, 4H), 2.09 (s, 4H). $^{19}$F NMR (500 MHz, DMSO-$d_6$) δ 66.85. LC-MS (ESI-API): calculated for $C_{26}H_{24}FN_4O_3S^+$ [M+H]$^+$: 491.2, found: 491.1.

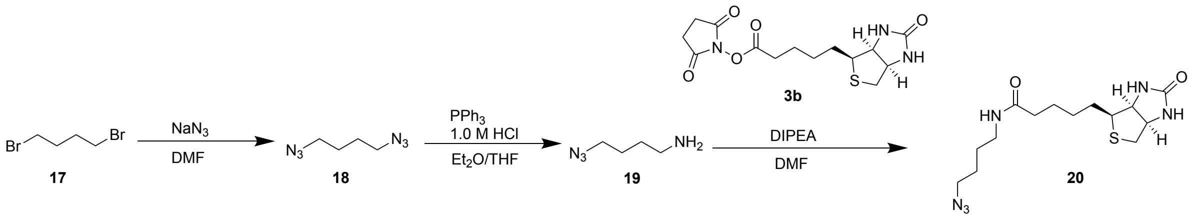

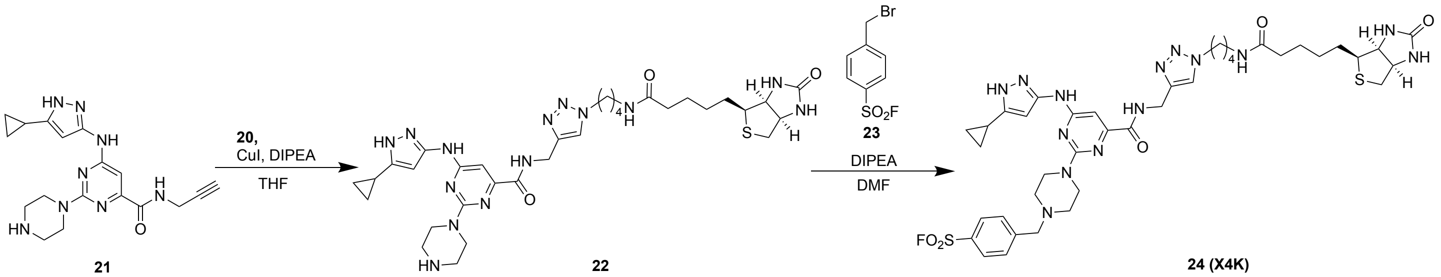

**Compound 18**: To a stirred solution of **17** (1.6 g, 7.5 mmol) in DMF (7.0 mL) was added $NaN_3$ (1.9 g, 29 mmol). The reaction was heated up to 80 °C and was allowed to stir at that temperature overnight before it was diluted with EtOAc (15 mL) after cooling to room temperature. The resultant mixture was washed with water (2 × 20 mL) and brine (2 × 20 mL), dried over anhydrous $Na_2SO_4$, and filtered. The solvent was evaporated under vacuum to give compound **18** (1.0 g, 95%) as a solid. $^1$H NMR (400 MHz, $CDCl_3$) δ 3.33 (tt, $J$ = 4.4, 1.9 Hz, 4H), 1.68 (tp, $J$ = 3.8, 1.1 Hz, 4H).

**Compound 20**: To a stirred solution of **19**[36] (0.12 g, 1.1 mmol) and **3b** (0.23 g, 0.67 mmol) in DMF (4.0 mL) was added DIPEA (0.26 g, 0.35 mL, 2.0 mmol). The reaction mixture was allowed to stir at room temperature overnight before it was quenched with water (15 mL). The resultant mixture was filtered and the solid was washed with water (2 × 15 mL) and brine (2 × 15 mL), dried over anhydrous $Na_2SO_4$, and filtered. The solvent was evaporated under vacuum to give compound **20** (0.19g, 85%) as a white solid. $^1$H NMR (400 MHz, DMSO-$d_6$) δ 7.79 (t, $J$ = 5.8 Hz, 1H), 6.42 (s, 1H), 6.35 (s, 1H), 4.31 (dd, $J$ = 7.7, 4.9 Hz, 1H), 4.17 – 4.07 (m, 1H), 3.17 – 2.98 (m, 3H), 2.83 (dd, $J$ = 12.4, 5.1 Hz, 1H), 2.58 (d, $J$ = 12.4 Hz, 2H), 2.05 (t, $J$ = 7.4 Hz, 2H), 1.92 (s, 1H), 1.74 – 1.21 (m, 10H).

**Compound 22**: To a stirred solution of **21**[37](54 mg, 0.15 mmol), **20** (50 mg, 0.15 mmol) and DIPEA (97 mg, 0.13 mL, 0.75 mmol) in THF (2.0 mL) were added CuI (14 mg, 75 $\mu$mol). The reaction mixture was allowed to stir at room temperature overnight. The crude product was purified by prep-HPLC with ACN:water (10:90 to 95:5) to give compound **22** (10 mg, 11%) as a white solid. $^1$H NMR (400 MHz, DMSO-$d_6$) δ 9.13 (t, *J* = 6.2 Hz, 1H), 8.79 (s, 1H), 7.91 (s, 1H), 7.78 (t, *J* = 5.7 Hz, 1H), 6.41 (s, 1H), 6.36 (s, 1H), 4.49 (d, *J* = 6.2 Hz, 1H), 4.31 (q, *J* = 6.3 Hz, 2H), 4.12 (dd, *J* = 7.9, 4.4 Hz, 1H), 4.00 (s, 3H), 3.18 (d, *J* = 8.1 Hz, 4H), 3.06 (dd, *J* = 15.2, 7.4 Hz, 3H), 2.82 (dd, *J* = 12.4, 5.0 Hz, 1H), 2.04 (t, *J* = 7.3 Hz, 3H), 1.90 (tt, *J* = 8.7, 5.0 Hz, 2H), 1.77 (t, *J* = 7.6 Hz, 2H), 1.67 – 1.06 (m, 14H), 0.93 (dt, *J* = 8.4, 3.2 Hz, 2H), 0.69 (dt, *J* = 6.5, 3.2 Hz, 2H).

**Compound 24 (X4K)**: To a stirred solution of **22** (11 mg, 16 $\mu$mol) and 4-(bromomethyl)benzenesulfonyl fluoride **23** (8.0 mg, 32 $\mu$mol) in DMF (0.50 mL) were added DIPEA (10 mg, 14 $\mu$L, 80 $\mu$mol). The reaction mixture was allowed to stir at room temperature for 1h before it was quenched with water (2.0 mL). The resultant mixture was filtered and the solid was washed with $Et_2O$ (2.0 mL) to give compound **24** (4 mg, 28%) in its TFA salt form as a white solid. $^1$H NMR (500 MHz, DMSO-$d_6$) δ 12.01 (s, 1H), 9.03 (s, 1H), 8.12 (s, 2H), 7.90 (s, 1H), 7.78 (s, 3H), 6.41 (s, 1H), 6.35 (s, 1H), 4.48 (d, *J* = 6.1 Hz, 2H), 4.31 (t, *J* = 7.6 Hz, 3H), 4.11 (s, 1H), 3.88 – 3.66 (m, 4H), 3.05 (dd, *J* = 13.7, 7.1 Hz, 4H), 2.81 (dd, *J* = 12.6, 5.2 Hz, 1H), 2.64 (s, 1H), 2.60 – 2.53 (m, 4H), 2.37 (s, 1H), 2.03 (t, *J* = 7.5 Hz, 2H), 1.87 (s, 1H), 1.76 (t, *J* = 7.7 Hz, 2H), 1.59 (s, 1H), 1.47 (d, *J* = 8.6 Hz, 4H), 1.38 – 1.21 (m, 6H), 0.90 (d, *J* = 8.1 Hz, 2H), 0.65 (s, 2H). $^{19}$F NMR (500 MHz, DMSO-$d_6$) δ 66.63. LC-MS (ESI-API): calculated for $C_{39}H_{52}FN_{14}O_5S_2^+$ [M+H]$^+$: 879.4, found: 879.3.

**NMR spectra of final compounds:**

**K60P- $^{1}H$ NMR (400 MHz, DMSO-$d_6$)**

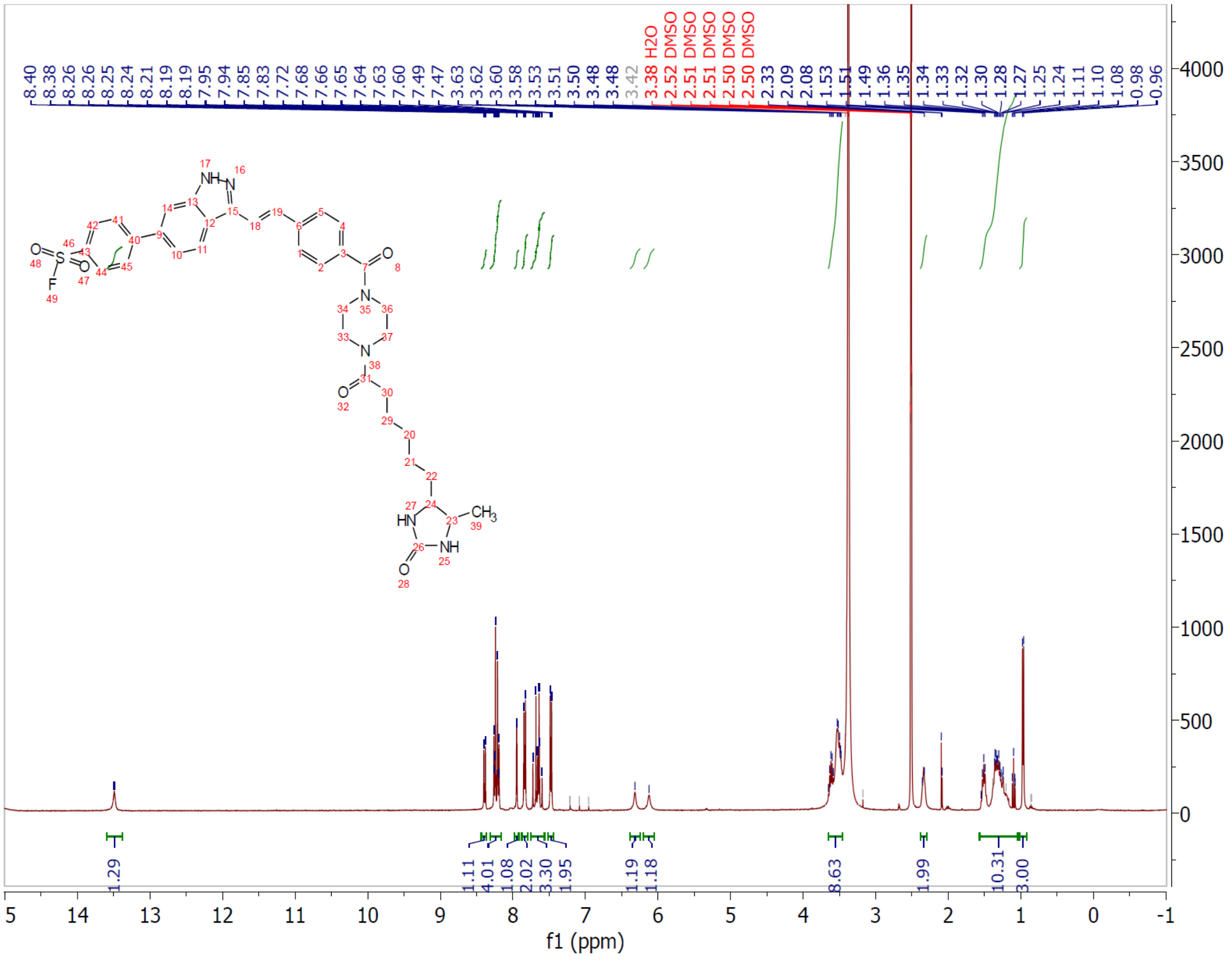

**K60P-Bio- $^{1}$H NMR (400 MHz, DMSO-$d_6$)**

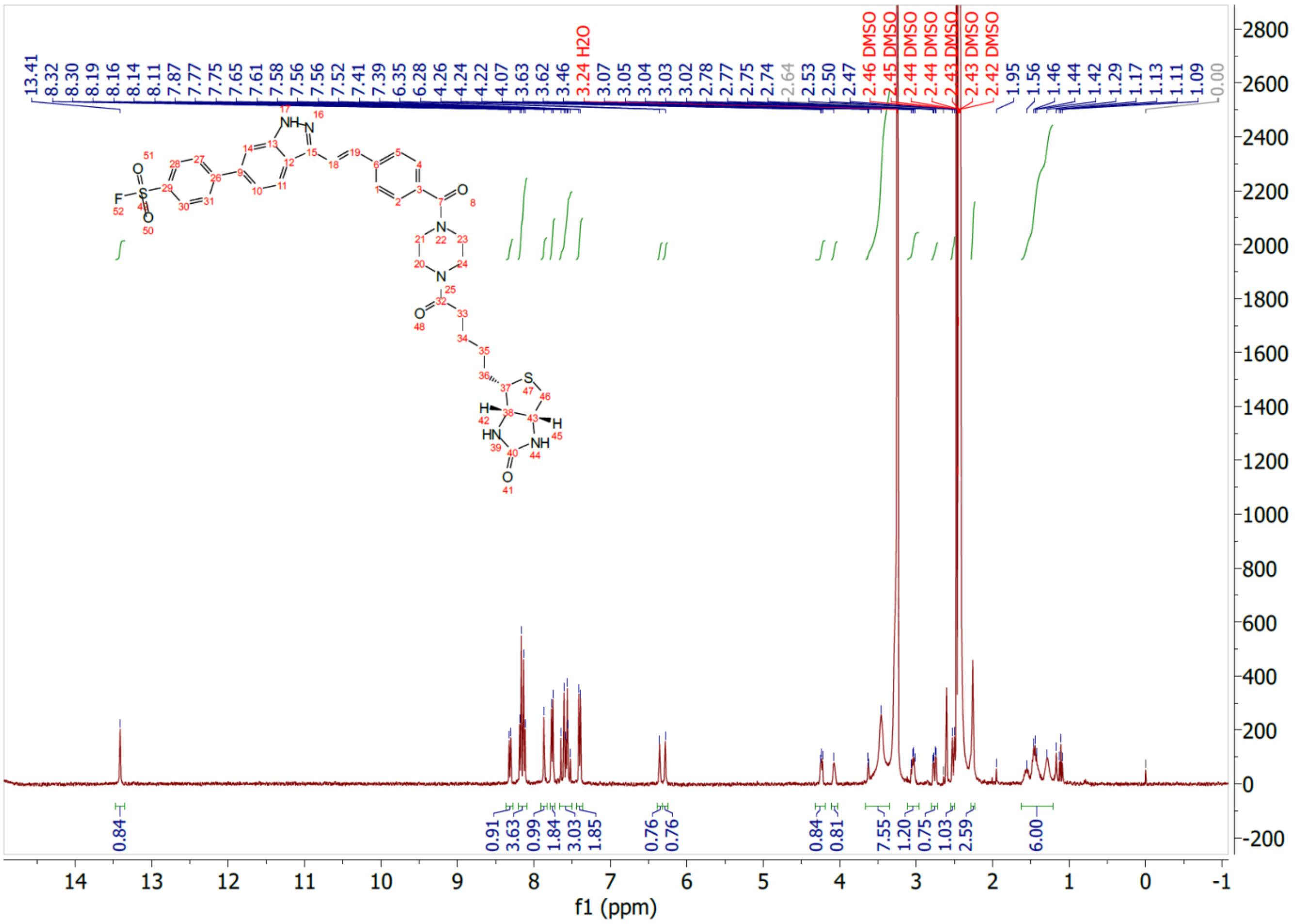

**PC-2-52- $^{1}$H NMR (400 MHz, DMSO-$d_6$)**

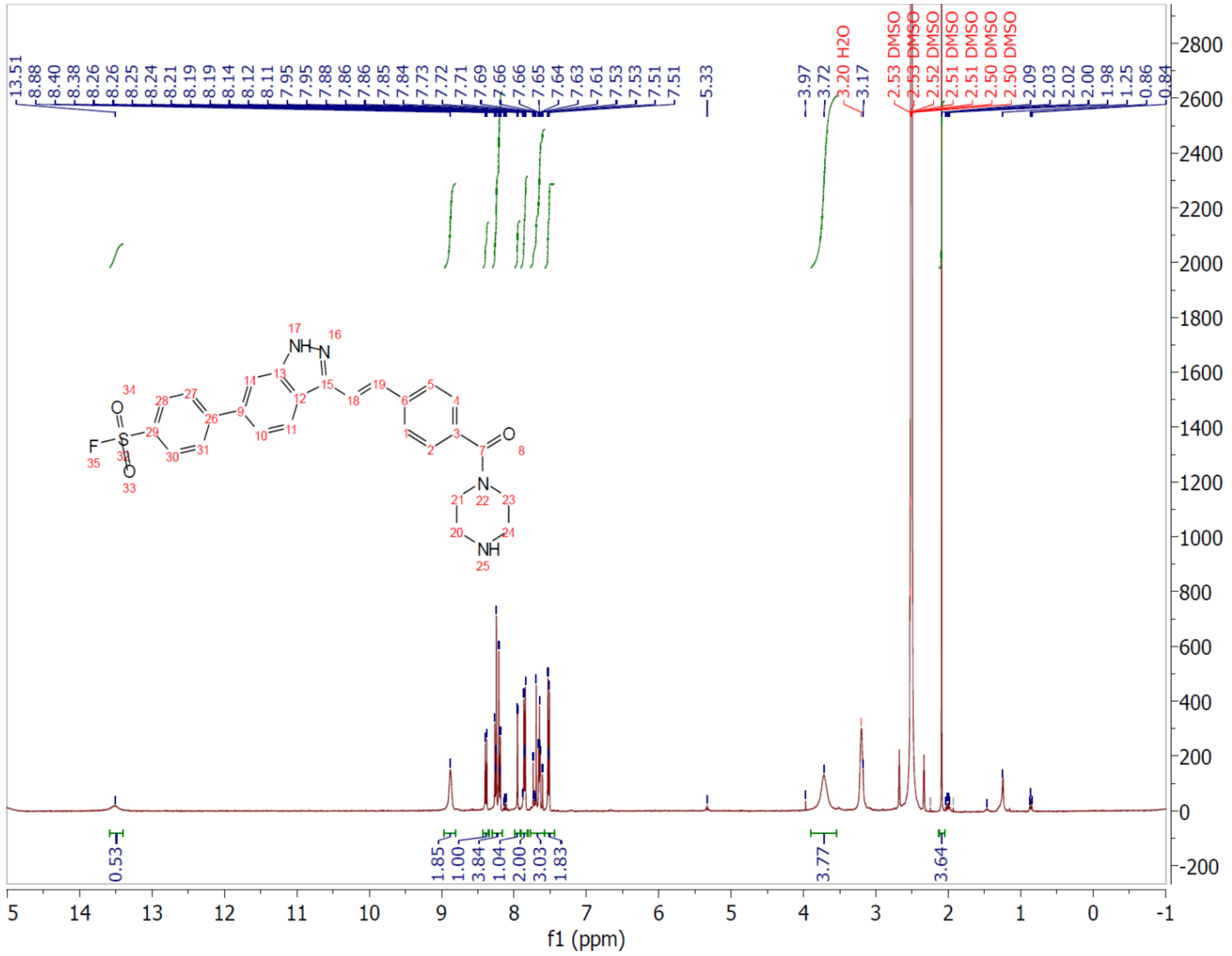

**K50P- [1]H NMR (400 MHz, DMSO-$d_6$)**

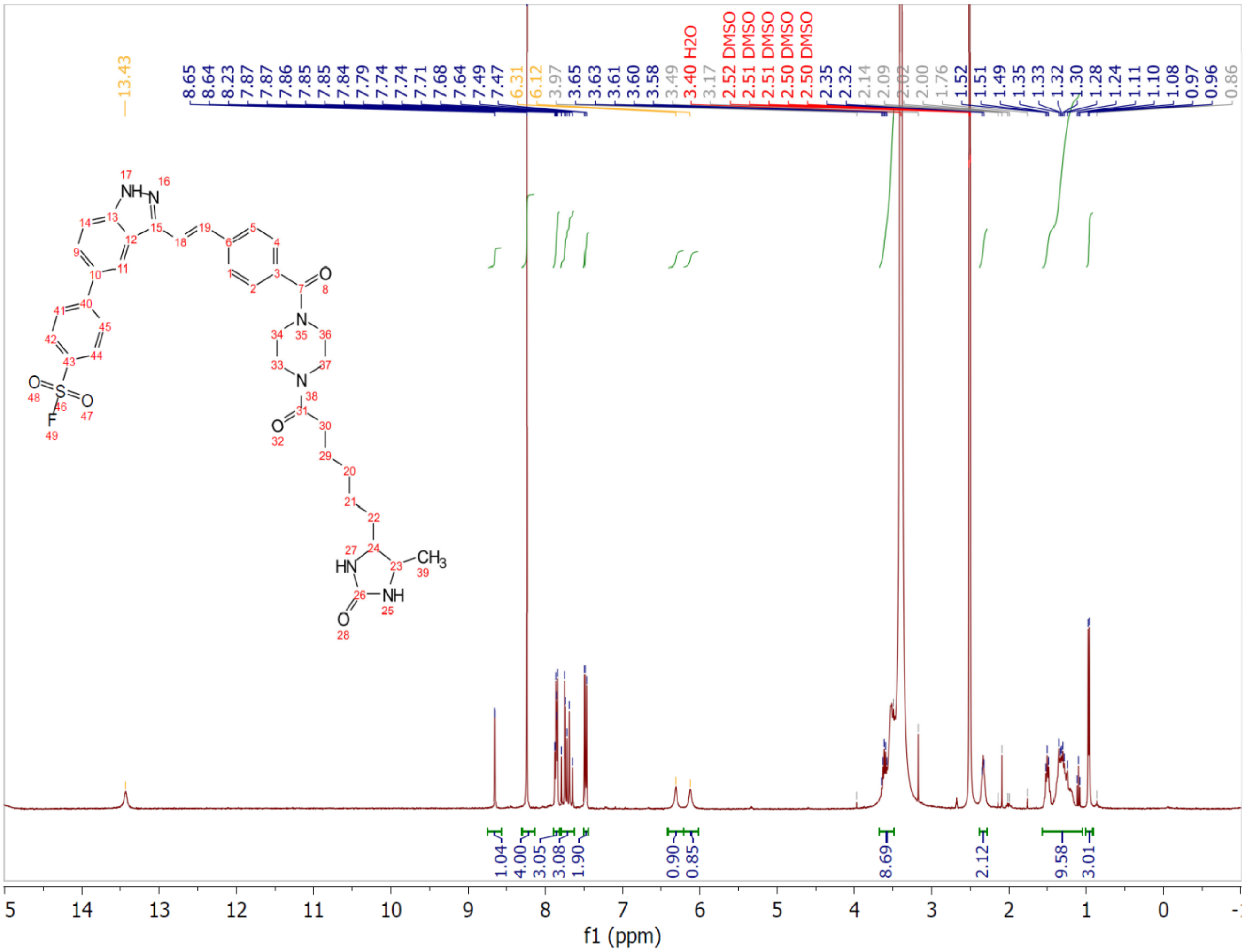

X4K- $^{1}$H NMR (500 MHz, DMSO-$d_6$)

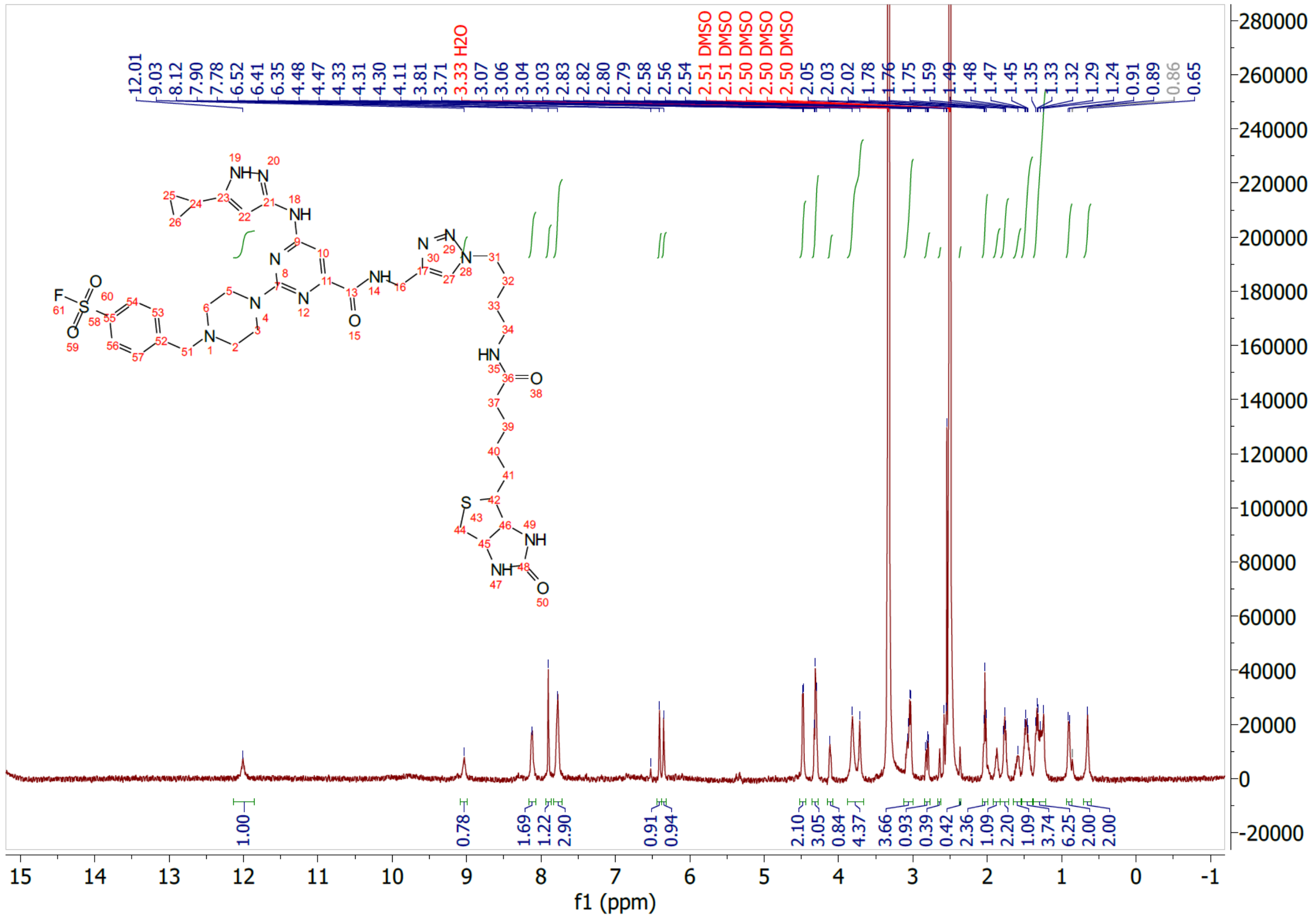
